# Experimental evolution in maize with replicated divergent selection identifies plant-height associated SNPs

**DOI:** 10.1101/2024.02.26.582128

**Authors:** Mila Tost, Cathy Westhues, Ginnie Morrison, Dietrich Kaufmann, Timothy Beissinger

**Affiliations:** Department of Crop Science, Division of Plant Breeding Methodology, University of Goettingen, Carl-Sprengel-Weg 1, Goettingen, 37075, Germany; Center for Integrated Breeding Research, University of Goettingen, Carl-Sprengel-Weg 1, Goettingen, 37075, Germany; University of Missouri, Division of Biological Sciences, 105 Tucker Hall, Columbia, MO, 65211, USA

**Keywords:** Experimental evolution, Population genetics, maize, selection signature mapping, plant height

## Abstract

Experimental evolution studies are common in agricultural research, where they are often deemed “long term selection”. These are often used to perform selection mapping, which involves identifying markers which were putatively under selection based on finding signals of selection left in the genome. A challenge of previous selection mapping studies, especially in agricultural research, has been the specification of robust significance thresholds. This is in large part because long term selection studies in crops have rarely included replication. Usually, significance thresholds in long term selection experiments are based on outliers from an empirical distribution. This approach is prone to missing true positives or including false positives. Under laboratory conditions with model species, replicated selection has been shown to be a powerful tool, especially for the specification of significance thresholds. Another challenge is that commonly used single-marker based statistics may identify neutral linked loci which have hitchhiked along with regions that are actually under selection. In this study, we conducted divergent, replicated selection for short and tall plant-height in a random mating maize population under real field conditions. Selection of the 5% tallest and shortest plants was conducted for three generations. Significance thresholds were specified using the false discovery rate for selection (FDRfS) based on a window-based statistic applied on a statistic leveraging replicated selection (*F_ST_Sum*). Overall, we found 3 significant regions putatively under selection. One region was located on chromosome 3 close to the plant-height genes *Dwarf1* and *iAA8*. We applied a haplotype block analysis to further dissect the pattern of selection in significant regions of the genome. We observed patterns of strong selection in the subpopulations selected for short plant height on chromosome 3.

## Introduction

The purpose of experimental evolution studies is to understand how populations evolve in response to a selective pressure (Janick, 2010a, 2010b). In agricultural research, they are commonly employed and often deemed “long term selection” studies (Dudley and Lambert, 2010; Moose, Dudley and Rocheford, 2004, Beissinger *et al.,* 2014). For instance, the Illinois long-term selection experiment for maize protein content began in 1896 at the University of Illinois and has been conducted ever since (Moose, Dudley and Rocheford, 2004; Dudley and Lambert, 2010). Experimental evolution studies are also common in model organisms like mice (Firman *et al*., 2015; Palma-Vera et al., 2022), *Drosophila melanogaster* (Phillips *et al*., 2016; Mallard, Afonso and Teotónio, 2023)*, E. coli* (Barrick *et al*., 2009; Card *et al*., 2021)*, Mimulus guttatus* (monkey flowers) (Angert, Bradshaw Jr and Schemske, 2008; Tusuubira and Kelly, 2024), and yeast (Payen *et al*., 2016; Linder *et al*., 2021). Early experimental evolution studies focused on the evolution of phenotypes alone, but now sequencing technologies allow researchers to study the effect of selection at the DNA level (Burke *et al.,* 2010; Parts *et al*., 2011; Orozco-terwengel *et al.,* 2012; Remolina *et al.,* 2012).

Many experimental evolution studies in model organisms work with replicates (Teotónio and Rose, 2000; Phillips *et al*., 2020; Mallard, Afonso and Teotónio, 2023). These studies are often referred to as Evolve & Resequence (E&R) experiments (Phillips *et al*., 2020). E&R experiments commonly employ pooled sequencing of the different subpopulations (Orozco-terWengel *et al.,* 2012; Long *et al*., 2015; Phillips *et al*., 2020). In agricultural species, experimental evolution studies that leverage replicated selection populations are exceedingly rare (Dudley and Lambert, 2010, Beissinger *et al*., 2014; Kumar *et al*., 2021). On the other hand, replication plays a very important role in E&R studies because it can help to identify putatively selected sites and differentiate those from sites caused by drift (Turner and Miller, 2012; Otte and Schlötterer, 2020; Linder *et al*., 2021; Lai *et al*., 2024). This assumes that selection is repeatable, while randomly acting forces like drift are not (Burke, Liti and Long, 2014; Phillips *et al*., 2020, Otte and Schlötterer, 2020; Schlötterer, 2023).

Genetic drift is a complicating factor in experimental evolution studies because it can be difficult to differentiate from selection (Krimbas and Tsakas, 1971; Turner *et al*., 2011; Hirsch *et al*., 2014). This is a recurring challenge in unreplicated selection mapping studies, because without replicating the evolutionary process, no amount of sampling can estimate the variability associated with genetic drift (Krimbas and Tsakas, 1971; Barghi, Hermisson and Schlötterer 2020, Kumar *et al*., 2021; Lai *et al*., 2024). The variability introduced by genetic drift depends on population size and the number of generations of selection (Baldwin-Brown, Long and Thornton, 2014; Kessner and Novembre, 2015; Barghi, Hermisson and Schlötterer, 2020). Replicated selection populations can help differentiate neutral loci subject to drift alone, from putatively selected loci (Kofler and Schlötterer, 2014; Barghi *et al.,* 2019; Linder *et al.,* 2021). Replicates can be analyzed to calculate a null expectation and to derive an approximation of the number of false positive observations in the experiment (Turner and Miller, 2012; Lai *et al*., 2024).

In previous experimental evolution studies on crops, significance thresholds for identifying signals of selection were frequently derived based on outlier quantiles of the empirical distribution (e.g. Wang *et al*., 2018; Ayalew *et al*., 2020, Semagn *et al*., 2021). This assumes that selection is a locus-specific force; putatively selected loci should be found at the tail of the empirical distribution (Akey *et al*., 2002; Akey, 2009). Drift is assumed to act uniformly genome-wide, so drifting loci dictate the bulk of the empirical distribution (Akey *et al*., 2002; Akey, 2009). However, it has been recognized for a very long time that outlier-based thresholds are arbitrary (Akey *et al*., 2002; Akey, 2009).

For this study, we performed replicated divergent selection for plant height on a random mating maize population for three generations. Plant height is an ideal model trait because it is highly heritable and highly polygenic (Pfeiffer *et al*., 2014, Mazaheri *et al.,* 2019). We replicated selection on four subpopulations, two of which were selected for tall plant height and two which were selected for short plant height. Each subpopulation was maintained with a population size of approximately 5,000 individuals. Every generation, we selected the 5% tallest or shortest plants (∼250 plants). The truncation threshold of 5% tallest or shortest plants was calculated based on 96 randomly sampled plants from each subpopulation (in total 384 plants). The 384 randomly sampled plants from the fourth generation were also used for genotyping-by-sequencing (GBS) to conduct selection mapping.

Regions corresponding to selection can be quite large and contain neutral linked loci (Tobler *et al*., 2014; Franssen, Barton and Schlötterer, 2016; Barghi and Schlötterer, 2019). Single marker-based statistics like *F_ST_* are prone to identify neutral loci that have hitchhiked along with those that are under selection (Tobler *et al*., 2014; Franssen, Barton and Schlötterer, 2016). Several experimental evolution studies have additionally investigated reconstructed haplotype blocks to disentangle the complex pattern of linkage (Michalak *et al*., 2019; Otte and Schlötterer, 2023; Chen, Pelizzola and Futschik, 2023). These investigations showed that often a large number of putatively significant sites corresponds to only a small number of significant haplotype blocks (Michalak *et al*., 2019; Chen, Pelizzola and Futschik, 2023). Therefore, we applied a cubic smoothing spline approach to identify window boundaries and corresponding values of a statistic denoted as “*Wstat”* for selection regions (Beissinger *et al*., 2015). This procedure may also reduce the inflation of observations due to linked loci (Barghi and Schlötterer, 2019; Chen, Pelizzola and Futschik, 2023). Finally, we generated significance thresholds based on Turner and Miller’s (2012) false discovery rate for selection (FDRfS), but instead of using single marker observations we calculated the FDRfS based on our previously defined regions. Finally, to assess the utility provided through replication, we compared these replication-based significance thresholds to those generated from the empirical distribution of *Wstat*. Additionally, we phased our data and investigated the haplotype blocks in the regions putatively under selection.

## Material and methods

### Plant materials and experimental design

The Shoepeg population is a maize Southern dent landrace population (Doebley *et al*., 1988), which was collected as open-pollinating variety in 1960 in Missouri, USA. We ordered Shoepeg material from the National Plant Germplasm System as accession PI 269743 (U.S. National Plant Germplasm System, 2021). Seeds were greenhouse-propagated to generate our starting population for selection. The formation of the Shoepeg experimental evolution population and results preceding the first generation of selection are described in Gyawali *et al*. (2019). To summarize, in 2016 the Shoepeg population was divided into four subpopulations, each consisting of approximately 5000 plants which were allowed to open-pollinate. In each subpopulation, plant height was measured (Figure 1). In total, 384 (96 x 4) individual plants out of approximately 20,000 (5000 x 4) plants were measured across all four subpopulations (Figure 1). Based on these plant height measurements, a truncation threshold corresponding to approximately the 5% shortest or tallest plants was calculated for each subpopulation (Figure 1) (Gyawali *et al*., 2019). Two subpopulations were selected for short plants, and two were selected for tall plants. The ∼5% shortest or tallest plants from each subpopulation were harvested as females to generate the next generation (Figure 1) (Gyawali *et al*., 2019).

**Figure 1:**
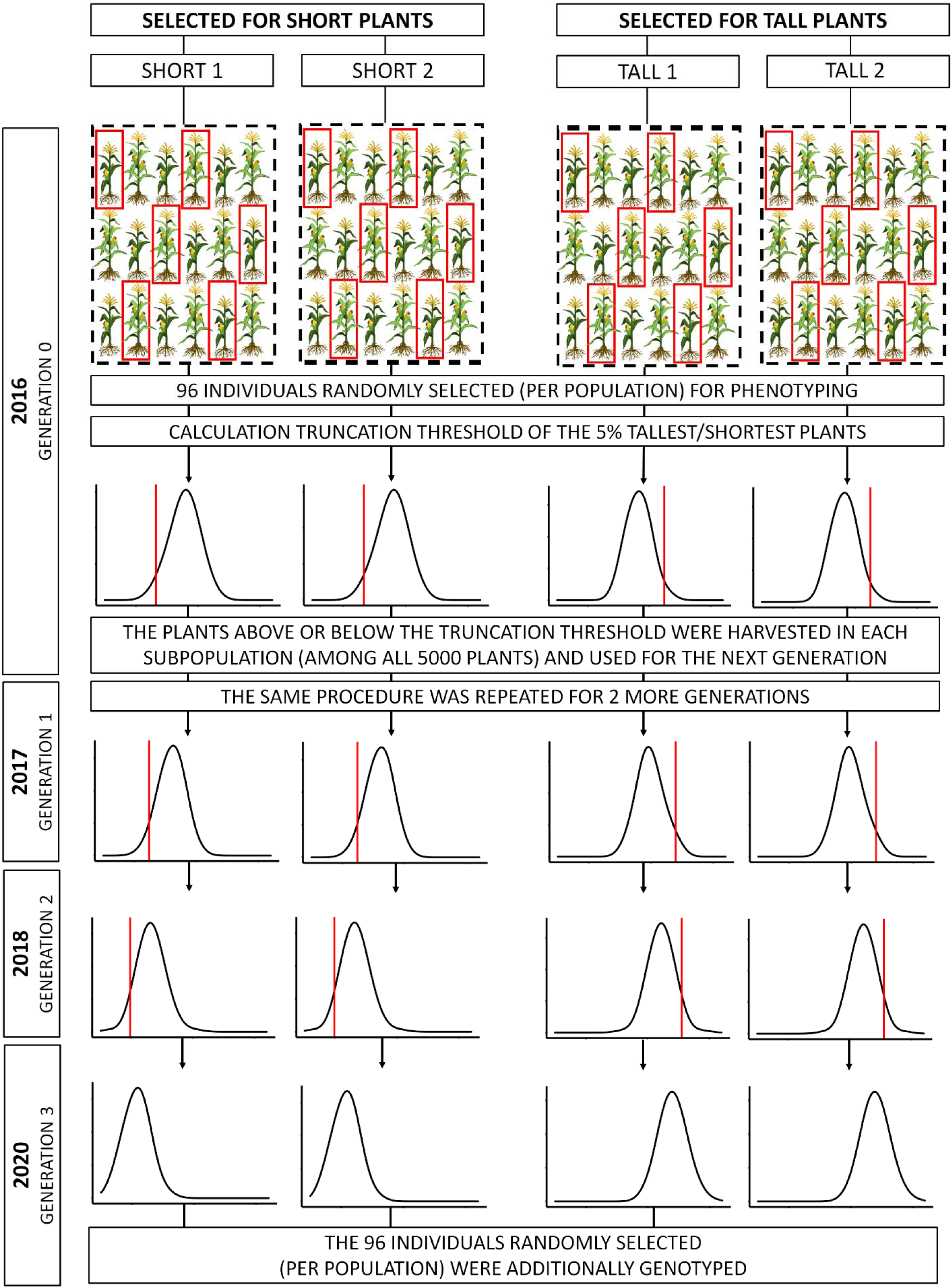
Simplified scheme for the development of subpopulations selected in divergent directions with their replicates for the three generations of selection. The subpopulations short 1 and 2 were selected for short plant height and the subpopulations tall 1 and 2 were selected for tall plant height.

This procedure was repeated in 2017 and 2018 (Figure 1). Ultimately, we generated four experimentally evolved subpopulations, two comprised of short-selected plants and two comprised of tall-selected plants (Figure 1). The populations were grown again in 2020 and plant height was measured for 96 randomly-chosen representative plants in each subpopulation (in total 384 plants). Additionally, tissue was collected from the same 384 plants for genotyping (Figure 1).

Every year after the first, the subpopulations were grown in isolated fields at least 200 m apart and away from any other maize to avoid contamination via cross-pollination (Nieh *et al*., 2014; Gyawali *et al*., 2019). In 2016, 2017 and 2018, the subpopulations were grown in Columbia, Missouri, USA (Gyawali *et al*., 2019). In 2020 the subpopulations were grown in Göttingen, Germany. In Missouri, each field had an approximate size of 0.1 ha (Gyawali *et al*., 2019). The row-spacing was 91 cm and the space between the plants was 15 cm (Gyawali *et al*., 2019). In Göttingen, the field size for three of the fields was 0.25 ha. One field was only 0.125 ha due to fewer seeds caused by a lower-than-expected harvest from one of the subpopulations in 2018. The distance between the seeds in Göttingen was approximately 33.3 cm and the row-spacing was approximately 1 m. The fields in Goettingen experienced noteworthy but non-devastating crow damage.

Plant height measurements were taken after flowering, from the ground up to the ligule of the flag leaf. Phenotypic data was analyzed using R version 4.0.3 (R Core Team, 2021). For phenotypic data analysis, data from 2016, 2017, 2018 and 2020 was used. The realized heritability from the breeder‘s equation (h^2^_bs_) was calculated as 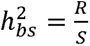 (Lush, 1937), where S is the selection differential and R is the response to selection. The selection differential was calculated as the difference between the mean of the population and the mean of the selected portion (Lush, 1937). The response to selection was calculated as difference between the mean of the previous generation and the mean of the next generation (Lush, 1937). This was done between all subsequent generations.

Flowering time was measured because in maize it tends to be correlated with plant height (Peiffer *et al*., 2014; Mazaheri *et al.,* 2019). This could lead to indirect selection for flowering time, thereby effecting the ability of plants to mate randomly. The possibility of this impacting the effective population size is addressed below.

### Genotyping, quality control and filtering

In 2020, the 96 randomly chosen plants from each subpopulation in generation four (384 individual plants in total) were genotyped using a genotyping-by-sequencing (GBS) approach. Leaf tissue from those 384 plants was freeze-dried and shipped to the University of Wisconsin Biotechnology center, Madison, WI, USA for DNA extraction, library preparation, and sequencing via one lane on a NovaSeq6000 machine. Each subpopulation was sequenced on its own sequencing run. Total sequencing yield from each run ranged from 125,000,000 to 300,000,000 fragments. Fragments had read lengths between 73 and 151 base pairs. GBS was conducted using paired-end sequencing with the restriction enzyme *Ape*KI (Elshire *et al*., 2011). For SNP variant calling the GB-eaSy pipeline was used (Wickland *et al*., 2017, Tange, 2011). Reads were aligned to the 5^th^ version of the B73 reference genome (Maize GDB, 2024) with the Burrows Wheeler Aligner (Li, 2013). In total, 15,390,958 SNPs were called by the GB-eaSy pipeline (Wickland *et al*., 2017). In the GB-eaSy pipeline, the SNPs were filtered for a minimum mapping and base quality of 30 with the BCFtools software (Danecek *et al*., 2021). After filtering, 8,302,778 SNPs were kept. The mapping quality ranged from 30 to 60, and was on average 57.44. The raw read depth ranged from 161 to 4,999, and was on average 1,705 reads. The average distance between adjacent markers was 257.67 bp.

Subsequent SNP filtering was conducted in R version 4.0.3 (R Core Team, 2021). Six individual samples with a marker coverage below 0.1 were removed from the analysis. The 10% quantile of marker coverage 0.6042, and 90% quantile was 0.7014 (supplementary figure S1). Markers with an average sample read depth below 1 or above 10 were removed. This filtering threshold was determined based on the distribution of read depth across all samples and markers (supplementary figure S2). Furthermore, we required that every SNP marker was observed at least 40 times in each subpopulation. Monomorphic and multi-allelic (>2) markers were removed from the data set. The final data set contained 4,029,092 SNP markers, which were used for the downstream analysis.

### Allele frequency estimation

The allele frequency at each SNP was calculated in R according to its maximum-likelihood estimate, which is computed by dividing the number of observations of a particular allele by the number of observations in total (Beissinger *et al*., 2014). The script is available at (https://github.com/milaleonie/ExpEvo_with_replicated_selection).

### Estimating effective population size

Effective population size was calculated using the known demographic parameters of the populations and based off the formula, 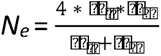, where *N_m_* is the number of male and *N_f_* the number of female individuals (Crow and Kimura, 1970). In every generation the ∼5% shortest or tallest plants were harvested, amounting to ∼250 plants. Under the assumption of complete random mating, all 5,000 plants could contribute as male parents for the next generation. In field trials, the assumption of random mating of all 5,000 plants is violated because of the effects of assortative mating (Allard, 1999). The number of male parents contributing to the next generation is reduced due to varying flowering dates and limited spatial pollen dispersion (Allard, 1999). Therefore, we evaluated the flowering dates of the 96 randomly chosen plants from each subpopulation to approximate the number of maximal simultaneously flowering tassels and silks in the entire population. We looked for the quantile in which the majority of silks is mature and receptive, because it was assumed that silks remain receptive up to five days after silk emergence (Nieh *et al*., 2014). We calculated the number of flowering tassels during this time interval and projected it onto the entire subpopulation. This procedure is explained in more detail in supplementary figure S3.

### Calculation of linkage disequilibrium (LD) decay

The extent of linkage disequilibrium (LD) was estimated based on 100,000 SNP markers with TASSEL v5 using the squared correlation between markers, r^2^, over a window size of 2,000 bp (Bradbury *et al*., 2007). We filtered the individuals out with a marker coverage below 0.6. For this analysis, we required that every SNP marker was observed at least 80 times in each subpopulation. This filtering resulted in 1,243,604 SNP markers. Additionally, we thinned the data set by randomly sampling 10,000 markers per chromosome. LD decay was modeled with a nonlinear regression model as expected value r^2^ under drift-recombination equilibrium as 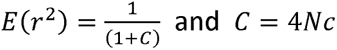 with N as the number of individuals at each site and C as the recombination coefficient between sites (Remington *et al*., 2001). We also created a pairwise LD heatmap with the R package LDheatmap for the regions which were putatively under selection (Shin *et al*., 2006).

### Scan for selection signatures

#### Calculation of *F_ST_Sum*

Our scan for selection was based on a statistic derived from *F_ST_*, which we call *<u>F_ST_Sum</u>*. First, *F_ST_* was calculated according to 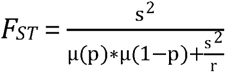, where s^2^ is the sample variance of the allele frequency between populations, µ(p) is the mean allele frequency between the subpopulations and r is the number of populations (Weir and Cockerham, 1984). The *F_ST_* statistic was calculated between the divergently selected subpopulations (Short 1 vs Tall 1; Short 2 vs Tall 2). Additionally, the *F_ST_* statistic was calculated between the subpopulations selected in the same direction (Short 1 vs Short 2; Tall 1 vs Tall 2). Next, the sum of *F_ST_* was calculated between the two non-redundant comparisons of divergently selected subpopulations (Short 1 vs Tall 1; Short 2 vs Tall 2) and subpopulation selected in the same direction (Short 1 vs Short 2; Tall 1 vs Tall 2). Assessing divergence among replicates can help in the distinction between putatively selected loci and false positives (Otte and Schlötterer, 2020; Chen, Pelizzola and Futschik, 2023).

#### Window-based analysis

We implemented a window-based analysis to assess linked selection. A cubic smoothing spline was applied to the single-marker based *F_ST_Sum* values. From the fitted spline, inflection points were calculated and used as window boundaries for newly defined regions. This analysis was performed with the R package GenWin (Beissinger *et al*., 2015). Briefly, the GenWin package returns window boundaries, which we used to define regions with linked markers that may be analyzed together (Beissinger *et al*., 2015). However, the windows are of variable sizes, so we used the W statistic (*Wstat*), as defined by Beissinger *et al*. (2015), to correct for this. *Wstat* is defined according to 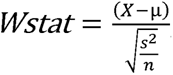, where X is the mean *F_ST_Sum* of a window, *µ* is the mean *F_ST_Sum* of the entire data set, *s^2^* is the sample variance of *F_ST_Sum* values across the entire data set and *n* is the number of markers. *Wstat* can therefore be used to compare regions, despite each consisting of a different number of markers (Beissinger *et al*., 2015).

#### Calculation of significance thresholds

Significance thresholds for selection were calculated in two ways: 1) based on the empirical distribution (as a baseline to compare to); and 2) based on the FDRfS (Turner and Miller, 2012).

1. The significance thresholds based on the empirical distribution were calculated by taking the 99.9^th^ and the 99.99^th^ percentile of the entire distribution of the *WStat* values (Akey, 2009; Beissinger *et al*., 2014; Hirsch *et al*., 2014; Ma *et al*., 2015; Cesconeto *et al*., 2017; Wang *et al*., 2018; Ayalew *et al*., 2020).
2. High *Wstat* values based on *F_ST_Sum* values calculated between divergently selected subpopulations are observed when the sites located in the region showed the same pattern of differentiation in both comparisons. Regions with high *Wstat* based on *F_ST_Sum* values calculated between subpopulations selected in the same direction are false positive observations for selection for plant height. By dividing the number of regions diverged between the subpopulations selected in the same direction by the number of regions diverged between the divergently selected subpopulation for each observation of *Wstat*, we obtained the false discovery rate for selection (FDRfS) for *Wstat*. The FDRfS was calculated for all observed windows (Turner and Miller, 2012). We identified the maximum *Wstat* value observed from comparisons between the subpopulations selected in the same direction, and used this as our significance threshold.

To visualize the scope of selection and drift, Turner and Miller (2012) plotted the allele frequency differences from the comparisons of divergently selected subpopulations and subpopulation selected in the same direction against each other. This plot is referred as the “Sauron plot”. We created this plot with *F_ST_*, which resulted in a “Quarter-Sauron” plot, because *F_ST_* ranges from 0 to 1, whereas allele frequency differences range from -1 to 1. Both plots are available in supplemental file S4. These plots depict how the FDRfS is computed and provides a visualization of the scope of drift and selection.

#### Haplotype estimation and haplotype block calculation

The data was phased using fastPHASE version 1.4.8 (Sheet and Stephens, 2006). We phased the genomic data of the regions significantly under selection with 10 iterations of the expectation-maximization (EM) algorithm (Sheet and Stephens, 2006). The phased haplotypes were used for plotting and the construction of the LD heatmap. Additionally, we phased 39 random regions across the entire genome, with the same physical length and marker density like the region significantly under selection from 9.411 to 10.423 Mb on chromosome 3. On all these regions we estimated haplotype blocks with the R package HaploBlocker (Pook *et al*., 2019). In the HaploBlocker package, a haplotype block is as a sequence of genetic markers of arbitrary length with a particular frequency in the population (Pook *et al*., 2019). The haplotype blocks only contain haplotypes with a similar sequence of markers (Pook *et al*., 2019). This helps to simplify the common haplotypes. The average number of haplotype blocks, the average block length and the total block number was calculated for all regions in the different subpopulations.

## Results

### Phenotypic evaluation

We observed notable and significant plant height differences between the subpopulations selected in opposite directions (Figure 2). After three generations of selection, the mean plant height difference between subpopulations selected in the opposite direction was 66.55 cm (P ∼ 0). The mean plant height decreased in the subpopulations selected for short plant height by 22.92 cm (P ∼ 0) and increased in the subpopulations selected for tall plant height by 30.61 cm (P ∼ 0). Throughout the experiment, the response to selection was always negative in the subpopulations selected for short plants and always positive in the subpopulations selected for tall plants (Figure 2).

**Figure 2:**
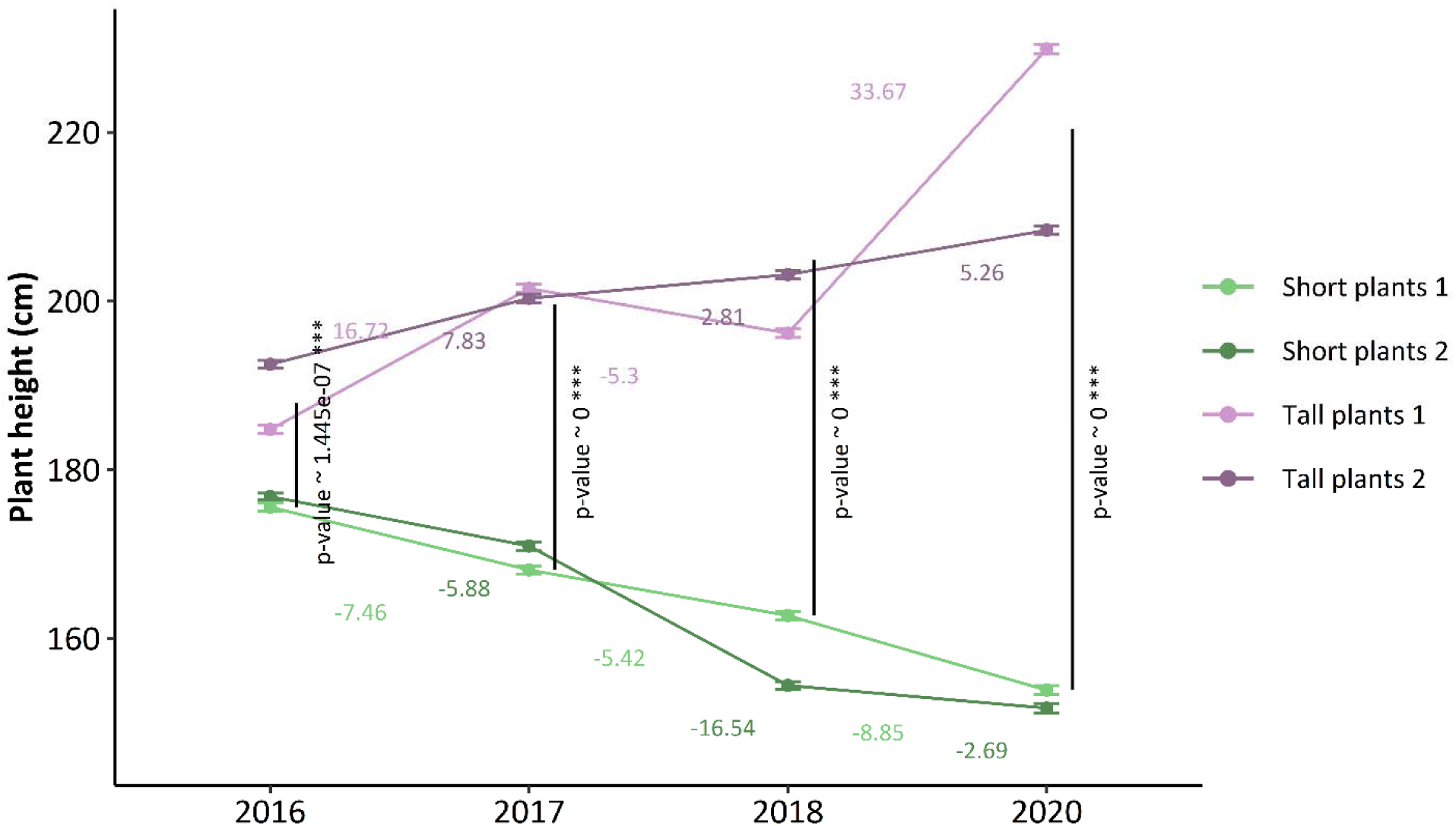
Mean plant height in each generation of selection. Error bars indicate a confidence interval for our estimation of means.

Realized heritability was computed by using the breeder‘s equation (h^2^_bs_) (Lush, 1937), and was estimated to be 0.3. Since each population was evaluated in a different environment (i.e. the year and location where it was selected), this application of the breeder’s equation should be considered only approximate. Plant height and flowering time were correlated in the subpopulations short 1, short 2, tall 1, and tall 2 with correlation coefficients of 0.2183 (P ∼ 0.0376), -0.0384 (P ∼ 0.7165), 0.2709 (P ∼ 0.0107) and 0.4572 (P ∼ 0), respectively.

### Effective population size and inbreeding

The effective population size under the assumption of complete random mating was 952 individuals. Due to differences in flowering time between plants, the assumption of random mating is violated. Based on observed flowering times, we estimated that ∼ 59.38% and ∼ 82.29% in the short and ∼ 56.25% and ∼ 38.54% in the tall subpopulations had the potential to pollinate one another. When we re-estimated effective population size based on this number of male plants, the effective population size was estimated ∼ 922 and ∼ 943 individuals in the short subpopulations and ∼ 918 and ∼ 885 individuals in the tall subpopulations.

### Comparison of significance thresholds and candidate gene identification

As a baseline for comparison, we also evaluated using outlier-based significance thresholds. Thresholds based on the 99.9^th^ and the 99.99^th^ percentile of the empirical distribution were exceeded, respectively, by 93 and 10 different regions (Figure 3A). Significance thresholds based on the FDRfS were exceeded by three different regions (Figure 3A). This demonstrates that had we utilized outlier-based significance thresholds in this experiment, we would have identified several false positive candidates for selection. *Wstat* values calculated between the subpopulations selected in the same direction are depicted in Figure 3B.

**Figure 3:**
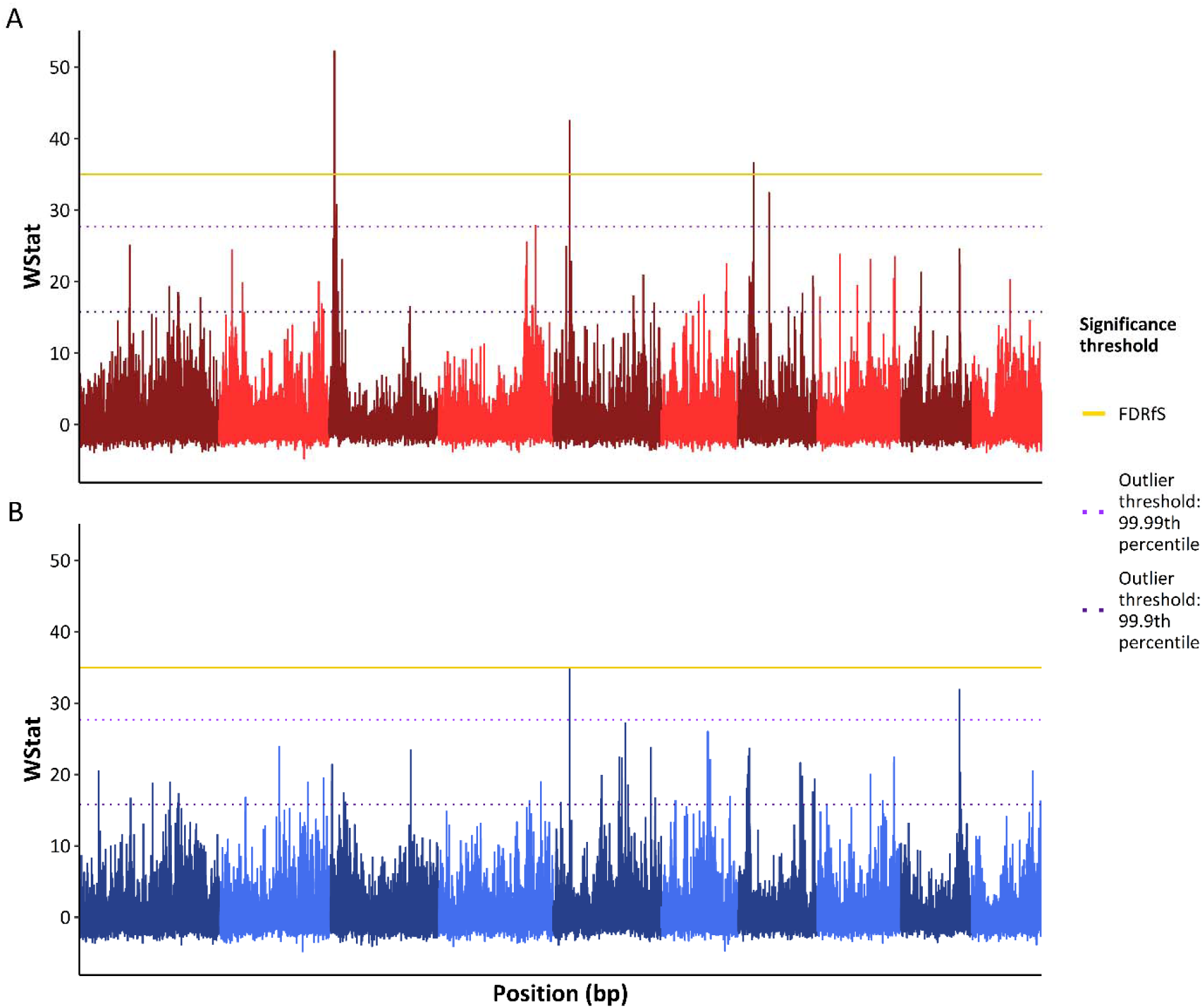
Differentiation along the entire genome expressed by *Wstat* calculated based on *F_ST_* at each SNP marker between the subpopulations selected in opposite directions (A) and the same direction (B) with the significance thresholds based on the 99.9^th^ percentile of the empirical distribution (light purple), based on the 99.99^th^ percentile of the empirical distribution (dark purple), and by the false discovery rate for selection (FDRfS) (yellow).

In supplemental S8, we included the results from *F_ST_Sum* values observed at single SNP markers plotted against their positions.

### Candidate plant height SNPs

All three regions under selection identified based on *Wstat* overlapped with coding regions in the B73 v5 annotated gene models (Maize GDB, 2024). The region with the highest *Wstat* value was located on chromosome 3 from 9.911 to 9.923 Mb (Figure 4). This 12 kb region is ∼ 138 kb away from *iAA8,* and is ∼ 518 kb away from *Dwarf 1* (Figure 4). The region is also 31 kb away from a significant marker identified in the Gyawali *et al*. (2019) study (Figure 4). The region was close to the coding region of other annotated gene models like *defective kernel5*, *glossy13* and *bzr6* (BZR-transcription factor 6) (Figure 4), but for these no previous evidence for their impact on plant height was found based on a literature review. We also observed other gene models, shown as yellow arrows, which are included in supplemental S10.

**Figure 4:**
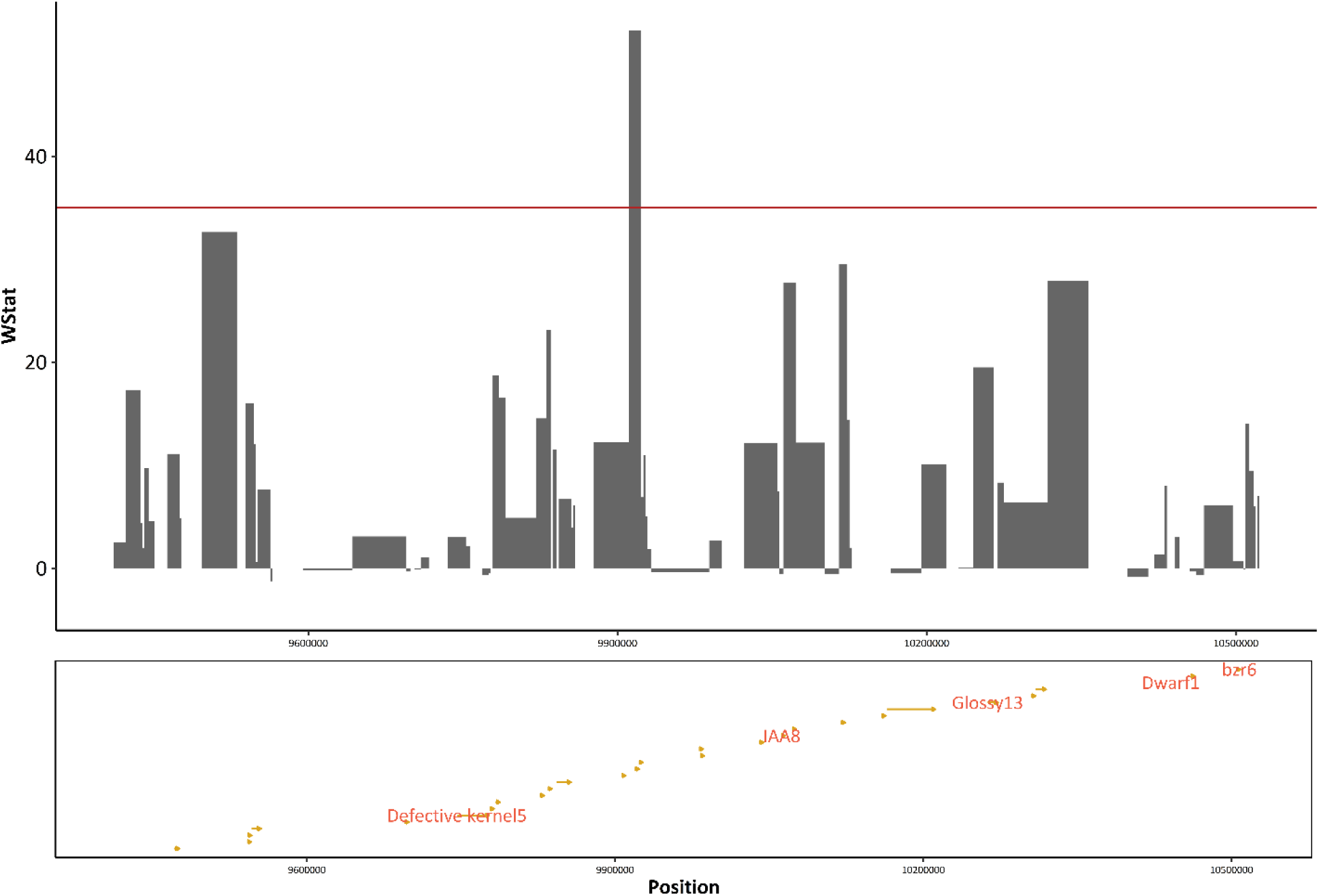
Putatively selected region on chromosome 3 from 9.911 to 9.923 Mb and 0.5 Mb up- and downstream and the coding regions of the gene models in this region.

The plant height genes *Dwarf1* and the *iAA8* have already been shown to be directly involved in plant height differences in many studies (Teng *et al*., 2013; Peiffer *et al*., 2014; Liu, Fernie and Yan, 2020; Ramos Báez *et al*., 2020). Therefore, these genes related to plant height were used as ground truth to test the method. Other neighboring gene models did not show a known connection to plant height based on a review of the literature.

The region on chromosome 5 from 30.103 to 30.115 Mb also had a length of 12 kb. This region was 32.39 kb downstream from the coding region of the gene *importin beta24* (Figure 5). The gene *importin beta24* is responsible for the mediation of the nucleocytoplasmic transport (Jin *et al*., 2022), and there is a study which reports a connection to dwarfism in Arabidopsis (Panda *et al*., 2020). However, a literature review revealed no evidence of its direct effects on plant height in maize. We also observed other gene models, shown as yellow arrows, which are included in supplemental S10.

**Figure 5:**
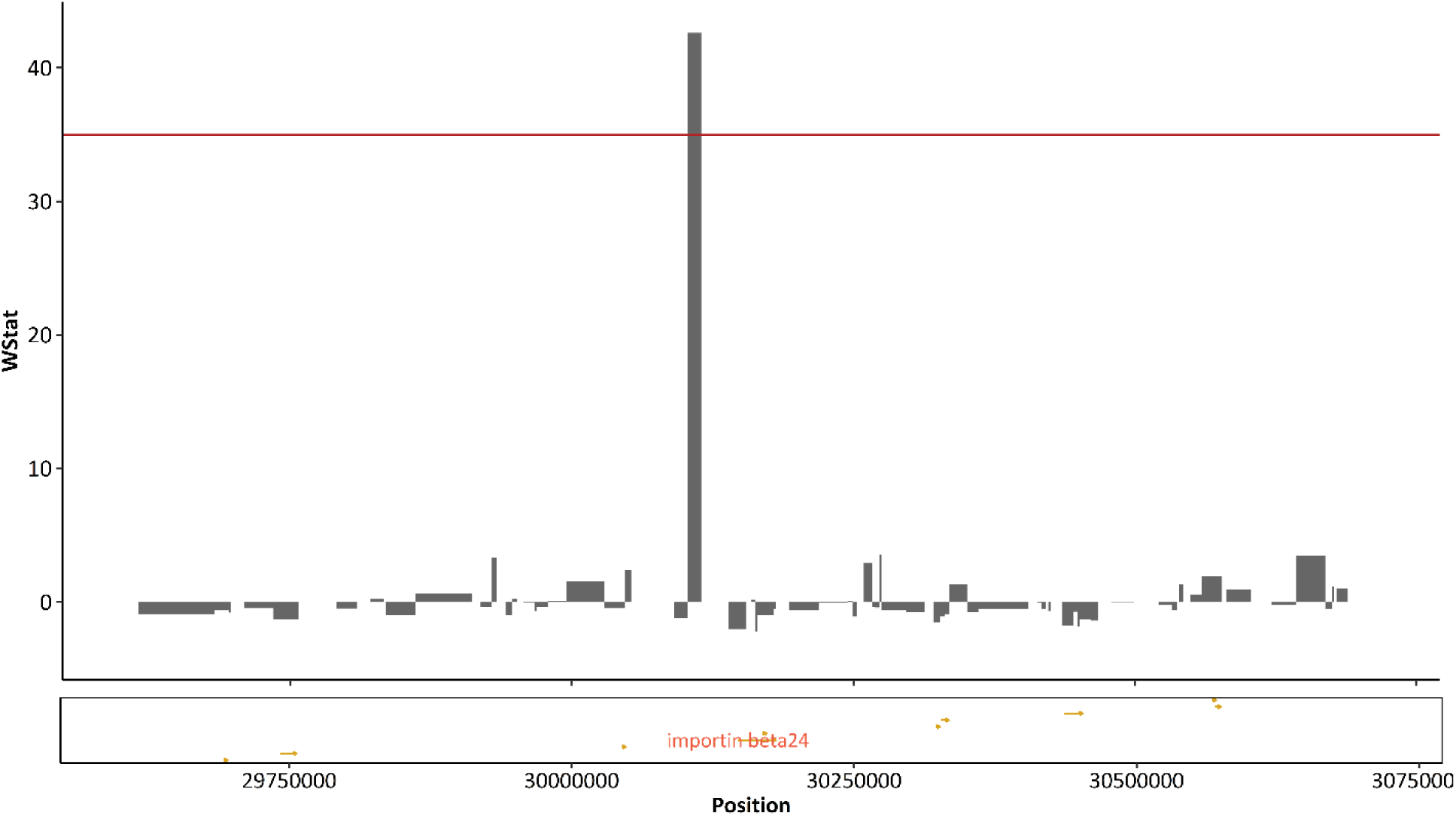
Putatively selected region on chromosome 5 from 30.103 to 30.115 Mb and 0.5 Mb upstream and downstream and the coding regions of the gene models in this region.

The region on chromosome 7 from 42.687 to 42.725 Mb had a length of 38 kb and was 435 kb upstream from the coding region of the gene model *beta expansin4* also referred as *expb4* (Figure 6). The gene *beta expansin4* is responsible for expansin protein which are capable of inducing cell wall extension in maize (Wu *et al*., 2001). But also, for this gene model no further evidence for their impact on plant height in maize was found based on a literature review. We also observed other gene models, shown as yellow arrows, which are included in supplemental S10.

**Figure 6:**
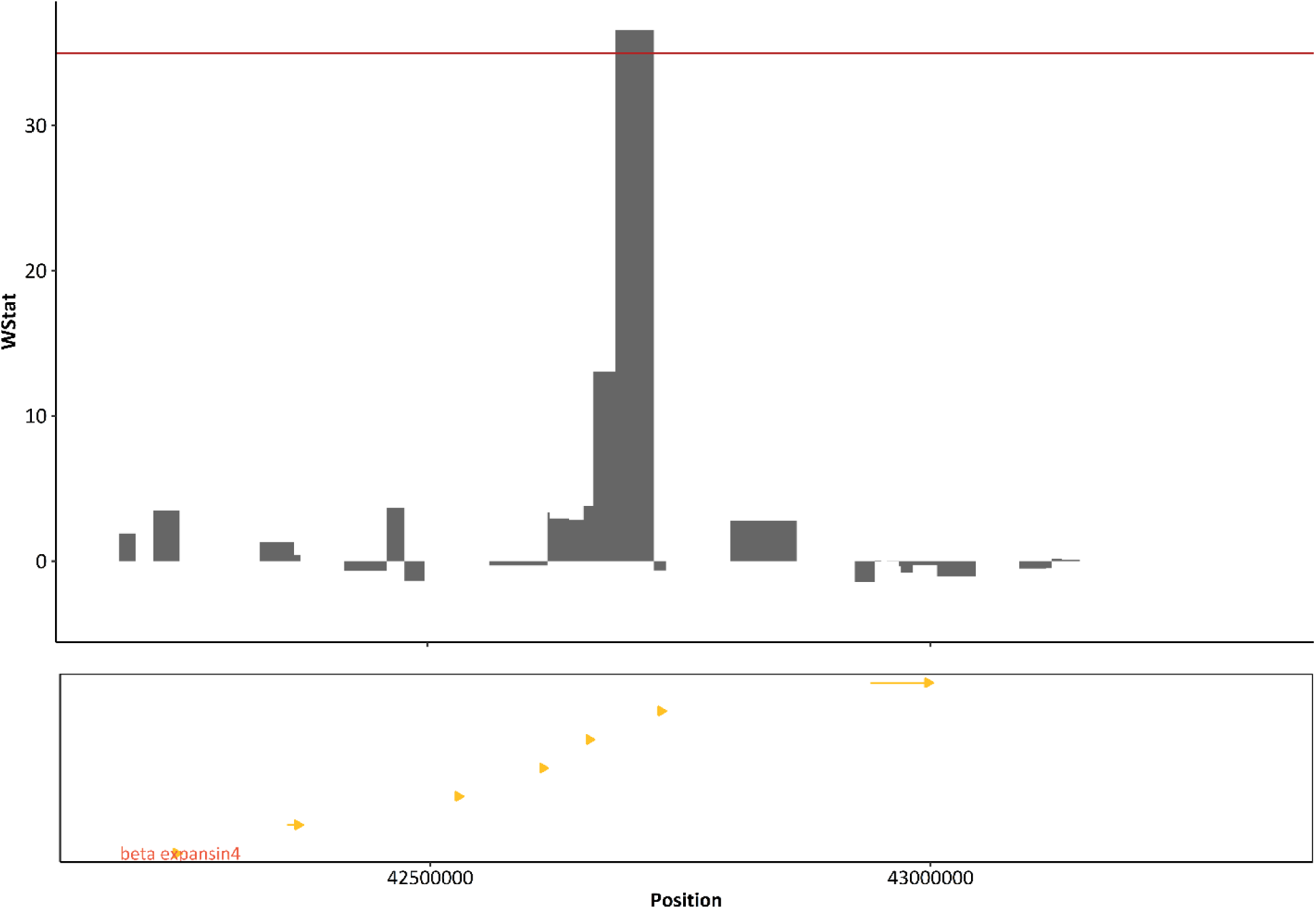
Putatively selected region on chromosome 7 from 42.687 to 42.725 Mb and 0.5 Mb up- and downstream and the coding regions of the gene models in this region.

### Haplotype block analysis

We investigated the regions putatively under selection in more detail by dissecting the haplotypes in the subpopulations. The haplotypes from the region from 9.911 to 9.923 Mb and 0.5 Mb up- and downstream on chromosome 3 are shown in Figure 7A. We observe different haplotypes across the entire region (Figure 7A). From ∼10.4131 to ∼10.4132 Mb, only 27.79 kb from the coding region of *Dwarf1*, we observed different alleles in the divergently selected subpopulations (Figure 7A). Also, from 10.0539 to 10.0646 Mb, located within the coding region of *iAA8*, a similar pattern of divergence is observed between the divergently selected subpopulations (Figure 7A). We also observed variation in haplotypes from 9.42 to 9.448 Mb and from 9.4712 to 9.563 Mb (Figure 7A). To supplement our haplotype investigation, we looked at LD across putatively selected regions in the different subpopulations. The pairwise LD heatmap indicates strong LD across the entire region from 9.911 to 9.923 Mb and 0.5 Mb up- and downstream in the subpopulations selected for short plant height (Figure 7B). We observed a low LD across the entire region in the subpopulations selected for tall plant height (Figure 7B).

**Figure 7:**
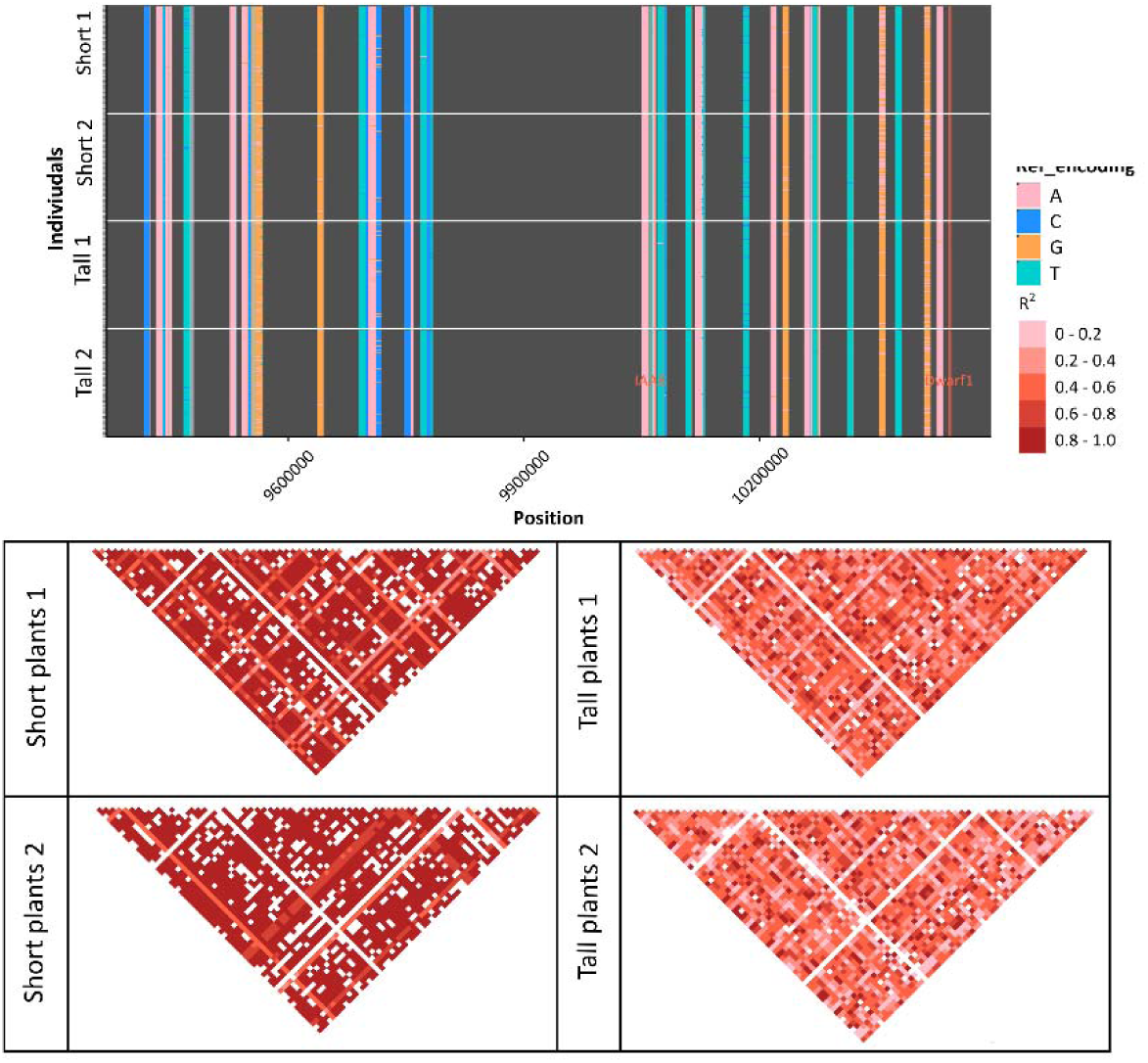
Putatively selected region on chromosome 3 from 9.911 to 9.923 Mb and 0.5 Mb up- and downstream and the observed haplotypes (A) and pairwise LD heatmap for the different subpopulations (B).

The haplotypes in the region from 30.103 to 30.115 Mb and the neighboring area of 0.5 Mb upstream and downstream on chromosome 5 are shown in Figure 8A. We observe large variation in the haplotypes around ∼ 29.8079, ∼ 29.8303, ∼ 30.2666 and ∼30.3253 Mb (Figure 9A). The pairwise LD heatmap indicates strong LD across the entire region in all subpopulations with no differences between the different subpopulations (Figure 8B).

**Figure 8:**
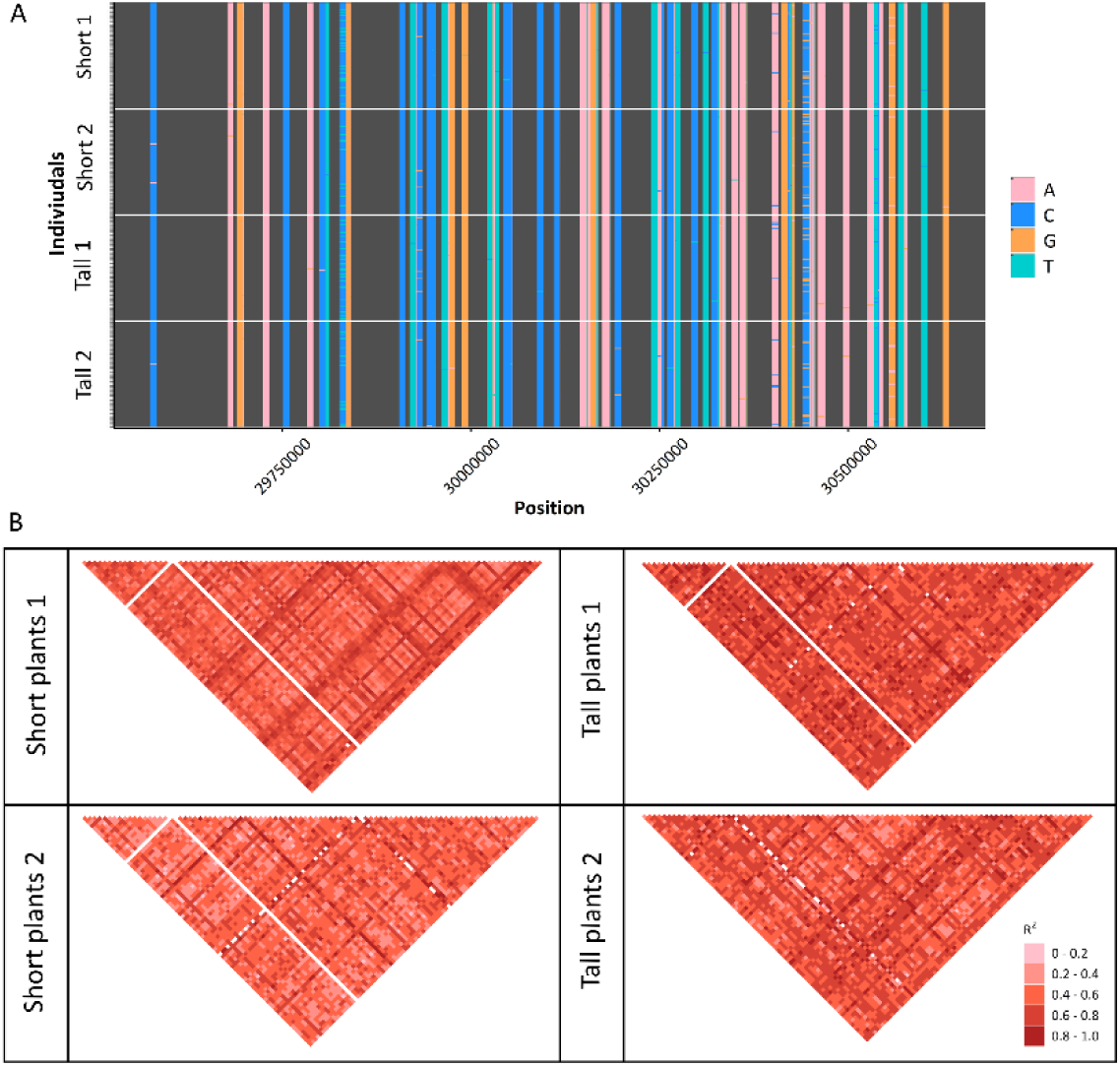
Putatively selected region on chromosome 5 from 30.103 to 30.115 Mb and 0.5 Mb upstream and downstream and the observed haplotypes (A) and pairwise LD heatmap for the different subpopulations (B).

**Figure 9:**
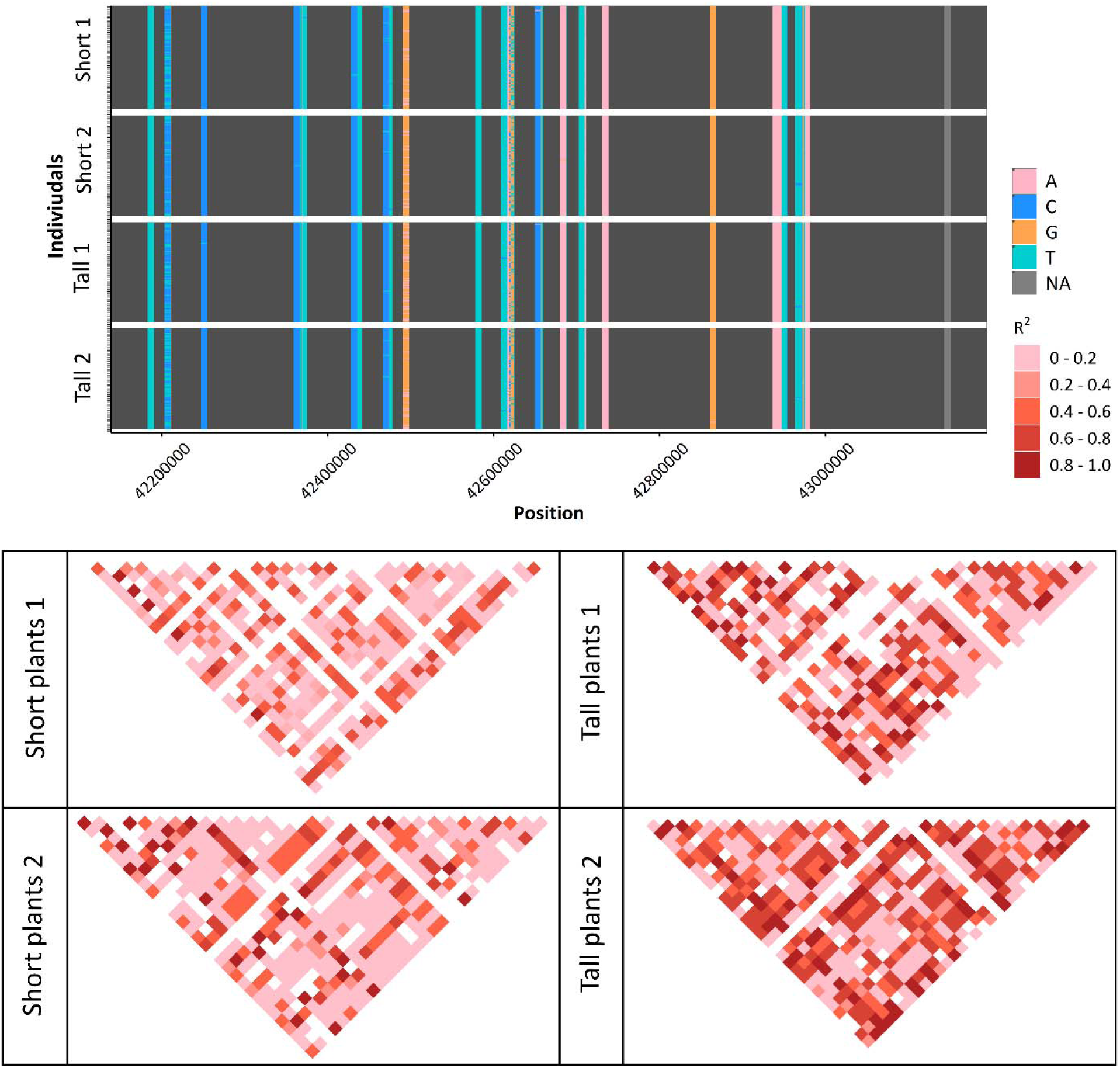
Putatively selected region on chromosome 7 from from 42.687 to 42.725 Mb and 0.5 Mb up- and downstream and the observed haplotypes (A) and pairwise LD heatmap for the different subpopulations (B).

Figure 9A shows the haplotypes in the region on chromosome 7 from from 42.687 to 42.725 Mb and 0.5 Mb up- and downstream. We observe large variation in the haplotypes around ∼ 42.6 Mb in many small regions (Figure 9A). We also observe variation around 42.2 Mb and from 42.432 to 42.49 Mb (Figure 9A). The pairwise LD heatmap indicates varying LD across the entire region in all subpopulations (Figure 8B). But overall, we observed a lower LD across the entire region in the subpopulations selected for short plant height and a stronger LD in the subpopulations selected for tall plant height (Figure 7B).

## Discussion

In contrast to most agriculture long-term selection experiments, our study included replication of the evolutionary process. This allowed us to directly compare the commonly used approaches to significance threshold specification leveraging replications. Our results demonstrate that significance thresholds specified using evolutionary replicates can widely vary from thresholds based on empirical outliers. Because linkage leads to selection operating on regions instead of individual markers, we used the R package GenWin (Beissinger *et al*., 2015) to calculate *Wstat* values based on a cubic smoothing spline applied to *F_ST_Sum*.

Significance thresholds based on the FDRfS identified fewer markers than outlier-based significance thresholds. Because these thresholds are based on replication, markers exceeding them are expected to correspond to true signals of selection. Previous studies applied significance thresholds based on the FDRfS on statistics like Chi-Square (Linder *et al*., 2021) and allele frequency differences (Turner and Miller, 2012). We adapted the calculation of the FDRfS to *F_ST_Sum*, a version of *F_ST_* which leverages replications. For comparison, we also explored calculating FDRfS from allele frequency differences (DiffStat) as was done by Turner and Miller (2012). The results are available in supplementary figures S4, S5 and S7. We identified more significant markers using the FDRfS from *F_ST_Sum* approach in comparison to allele frequency differences (DiffStat). This was also observed by Kofler and Schlötterer (2014). Our window-based approach seems to be the more suitable method than single marker based *F_ST_Sum* values since the single marker based *F_ST_Sum* would have returned many single markers tagging the same region as it is shown in supplemental file S9.

The region on chromosome 3 from 9.911 to 9.923 Mb was very close to a marker from previous study by Gyawali *et al*. (2019). Furthermore, the gene models *iAA8, Dwarf1*, *defective kernel5*, *glossy13* and *bzr6* were found close to this region. The plant height genes *iAA8* and *Dwarf1* are∼ 138 kb and ∼ 518 kb away from the putatively selected region from 9.911 to 9.923 Mb on chromosome 3.

*Dwarf1* is involved in the expression of gibberellin 3-oxidase (Teng *et al*., 2013). High expression of the gene leads to high levels of bioactive gibberellin and results in taller plants (Teng *et al*., 2013). When the *Dwarf1* gene is not expressed, plants exhibit the *Dwarf1* phenotype, which is characterized by dwarfism (Teng *et al*., 2013). Multiple gene expression studies have previously reported that this gene has a large effect on plant height (Teng *et al*., 2013; Liu, Fernie and Yan, 2020). In one experiment, this gene explained 32.3% of the phenotypic variance for plant height (Teng *et al*., 2013). The *iAA8* gene regulates auxin biosynthesis, transport, and signaling (Matthes *et al*., 2019; Ramos Báez *et al*., 2020). Auxin is a plant hormone and plays a key role in plant growth and development (Ramos Báez *et al*., 2020). The *iAA8* gene was also observed to regulate maize inflorescence architecture (Galli *et al*., 2015; Ramos Báez *et al*., 2020). Both genes appear to be causal for plant height differences based on the literature (Teng *et al*., 2013; Peiffer *et al*., 2014; Liu, Fernie and Yan, 2020; Ramos Báez *et al*., 2020). Therefore, these plant height genes were used as ground truth to test the method.

The *defective kernel 5* gene model was reported to cause reduced starchy endosperms and leads to a shrunken or collapsed endosperm (Zhang *et al*., 2019). *Glossy13* encodes a putative ABC transporter required for the accumulation of epicuticular waxes, which also help to control water loss (Li *et al*., 2013). The gene model *bzr6* is a transcription factor for brassinosteroids which are plant specific steroidal hormones (Manoli *et al*., 2017). Brassinosteroids contribute to regulating a broad spectrum of plant growth and developmental processes in maize (Manoli *et al*., 2017).

Very close to the coding region of *Dwarf1* and within the coding region *iAA8*, we observed different haplotypes in the divergently selected subpopulations (Figure 7A). Overall, we observed stronger LD in the putatively selected region on chromosome 3 in the subpopulations selected for short plants. These differences may indicate stronger selection in the subpopulations selected for short plants.

This could theoretically be a result of more inbreeding in the short subpopulations, which we tested by evaluating haplotype blocks in 39 random regions of the genome. The random regions were of equal size and marker density to the putatively selected region on chromosome 3. We observe on average 31 blocks in short 1, 30 blocks in short 2, 31 blocks in tall 1 and 34 blocks in tall 2. The blocks have an average block size of 402.8 bp in short 1, 427.36 bp in short 2, 409.01 bp in tall 1 and 404.76 bp in tall 2. This indicates that the tall subpopulations show similar levels of inbreeding to the short subpopulations. Results from the calculated haplotype blocks from the 39 random regions are available in supplementary figure S11 and S12.

Overall, a slow decline in LD was observed with increasing physical distance across all chromosomes (Supplemental S13). R^2^ of 0.1 was observed at a distance of 2 Mb (Supplemental S13). But when we consider the subpopulations separately in the pairwise LD heatmap, we observe very different results (Figure 7B). In the pairwise LD heatmap calculated for each subpopulation and the different regions, we observed strong variation in LD between the divergently selected subpopulations and between the different regions (Figure 7B).

The observation of strong LD in the putatively selected region on chromosome 3 in the short plant populations supports the argument that strong selection may act on the entire candidate gene region in theses populations (Franssen, Barton and Schlötterer, 2016; Barghi and Schlötterer, 2019, Otte and Schlötterer, 2020; Schlötterer, 2023). The strength of selection and the experimental design influence hitchhiking (Baldwin-Brown, Long and Thornton, 2014; Kofler and Schlötterer, 2014). We applied a strong selective regime in our experiment in comparison to previous studies (Kofler and Schlötterer, 2014; Long *et al*., 2015). Strong selection leads to more hitchhiking and a lower mapping resolution (Kofler and Schlötterer, 2014), which we observe in the short subpopulations. Another reason for strong selection acting on entire regions could be that many loci in this region are linked, resulting in joint selection (Schlötterer, 2023). However, strong LD may also persist due to more inbreeding in the short subpopulations. But we only observe strong LD in the short subpopulations in our putatively selected region on chromosome 3. On the other hand, we observed stronger LD in the tall subpopulations in our putatively selected region on chromosome 7 and similar LD patterns among all subpopulations in our putatively selected region on chromosome 5. When we calculated the pairwise LD heatmap for the different subpopulations in two random regions (supplementary figures S14 and S15), we did not observe a similarly strong difference in LD between the subpopulations as we did for the putatively selected region on chromosome 3 (Figure 7).

Even though we observed stronger LD in the tall subpopulations across our putatively selected region on chromosome 7, we also observed variation in LD. This may be because this small region is under selection. We also observed large variation in the haplotypes in this region, whereas most of the neighboring regions exhibit fixed haplotypes. The putatively selected region is only 22.78 kb from an uncharacterized protein (see supplemental S10), whereas *beta expansin4* is located 32.39 kb away from this region (Maize GDB, 2024). The gene *beta expansin4* is responsible for expansin protein which are capable of inducing cell wall extension in maize (Wu *et al*., 2001).

Close and within the other putatively selected regions on chromosome 5, several gene models were found but based on our literature review, no previous evidence for their impact on plant height was found. The gene model *importin beta24* is located on chromosome 5 and was 32.39 kb upstream from the putatively selected region on this chromosome (Maize GDB, 2024). Across this region and its neighboring area, we observed strong LD across all subpopulations. We observed variation in haplotypes in smaller neighboring segments of the coding region of *importin beta24*. This may indicate that this gene model is under selection. The gene *importin beta24* is responsible for the mediation of the nucleocytoplasmic transport (Jin *et al*., 2022), but there is a study which reports a connection to dwarfism in Arabidopsis (Panda *et al*., 2020). These gene models and the gene model *bzr6* on chromosome 3 may be interesting candidates for further studies to test their effect on plant height.

It is not only the strength of selection that can influence the number of variants selected in experimental evolution studies. The number of selected variants also depends highly on experimental design factors such as the population size, the duration of the experiment, the applied selective regime (Kofler and Schlötterer, 2014; Kessner and Novembre, 2015; Lai *et al*., 2024), and the species (Tobler *et al*., 2014; Barghi *et al*., 2017). Divergent selection can increase the power of the experiment in comparison to a single direction experiment (Kessner and Novembre, 2015). We observe differences regarding the resolution between the subpopulations selected in divergent directions. Even though the same strength of selection is applied in the divergently selected subpopulations, differences between the subpopulations can be observed (Moose, Dudley and Rocheford, 2004). But, these differences indicate that there is an increase in power due to divergent selection (Kessner and Novembre, 2015; Schlötterer, 2023).

Previous simulation studies investigating experimental evolution have demonstrated that replication is one of the most powerful tools to increase power (Kofler and Schlötterer, 2014; Otte & Schlötterer, 2020, Phillips *et al*., 2020). In fact, Kofler and Schlötterer (2014) observed that replication is more important for detecting variants than population size (Kofler and Schlötterer, 2014; Kessner and Novembre, 2015; Barghi, Hermisson and Schlötterer, 2020).

## Conclusion

In conclusion, our research demonstrates that replicated selection is a powerful tool in agricultural experimental evolution studies. We observed selection across replicates, even though we only selected for three generations. Moreover, we saw that a significance threshold based on the FDRfS is more reliable than existing methods for significance threshold identification. The application of a cubic smoothing spline to single marker-based statistics like *F_ST_Sum* to return window boundaries of regions which are putatively under selection helped collapse signal from multiple hitchhiking markers. Haplotype block analysis additionally helped to investigate the regions putatively under selection. We observe very different patterns of selection in divergently selected subpopulations, which might be due to selection. After more generations of selection, and a reduction of linkage between putatively selected loci and neutral loci, we may draw more certain conclusions about selection for the plant height genes. Furthermore, we might be able to identify significant markers close to our plant height candidate genes and novel gene loci without any associated gene models.

## Availability of data and materials

All the data and statistics about the present study has been included in the current manuscript in the form of figure and tables. Raw data are publicly available at figshare; https://figshare.com/projects/Shoepeg_ExpEvo2020/139981. Our analysis code is available publicly on github: https://github.com/milaleonie/ExpEvo_with_replicated_selection.

## Acknowledgements

Abiskar Gyawali contributed substantially to this study but passed away before submission. Journal policy prohibits him from being listed as a co-author because he could not approve the final manuscript. We consider Abi to be an important co-author of this manuscript, are grateful for his contributions, and miss him tremendously. Kate Guill was instrumental in conducting early-generation seed increase and field management in Missouri.

## Supplementary files

**S1:**
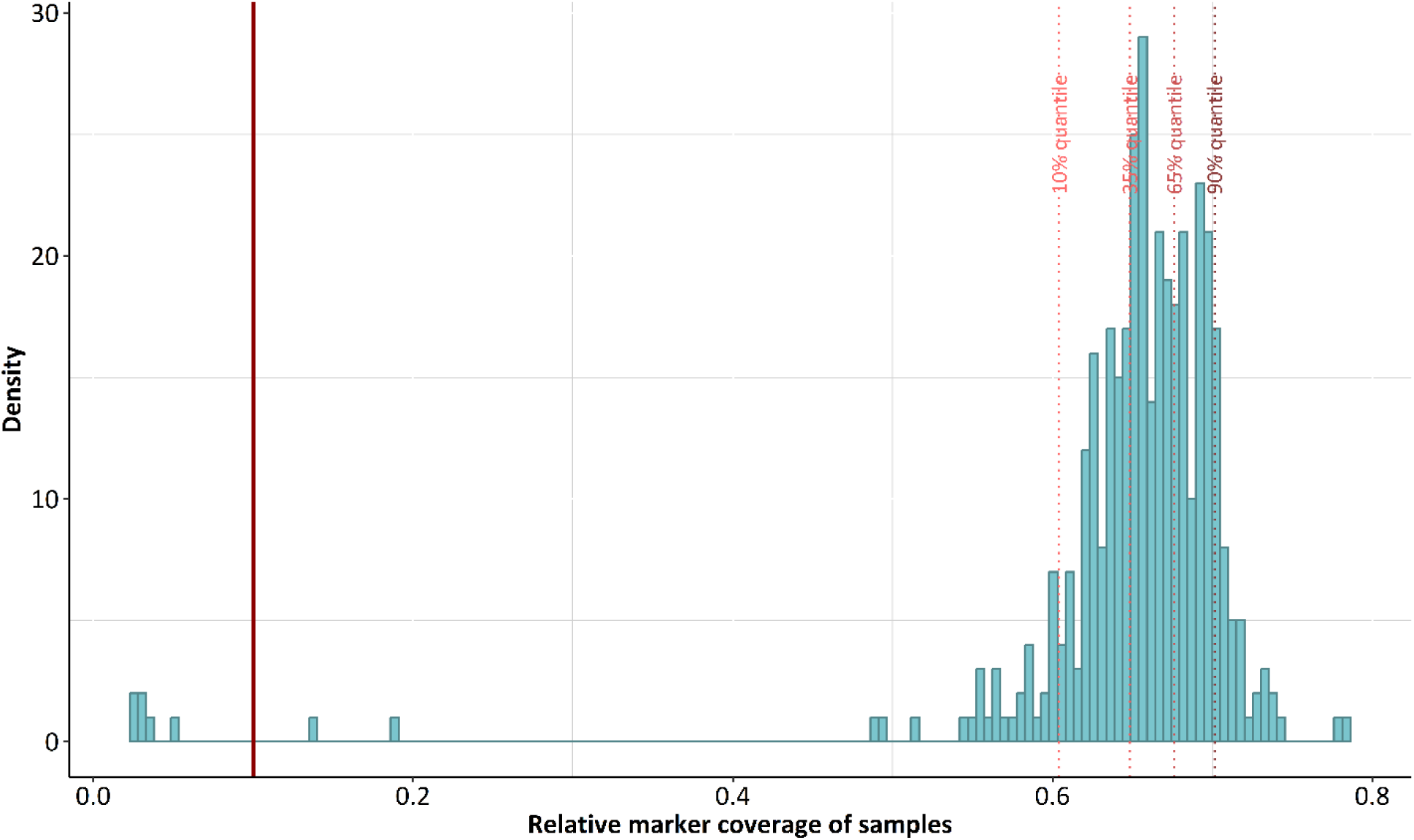
Distribution of marker coverage across samples with the 10%, 35%, 65% and 90% quantiles and the cut-off at a marker coverage of 0.1 (dark red).

**S2:**
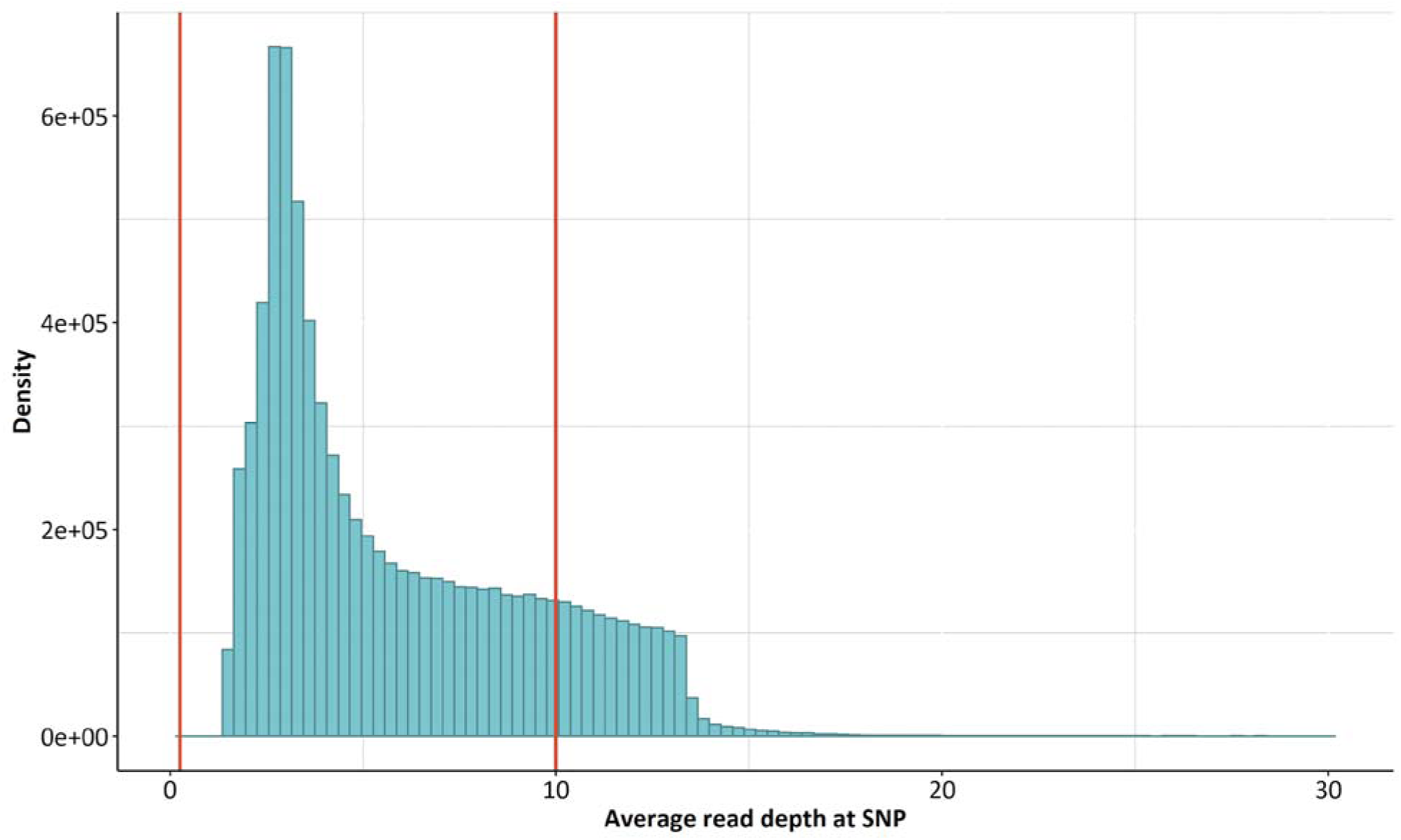
Distribution of read depth across all samples and markers. Red bars represent the cutoff threshold for markers with an average sample read depth below 1 or above 10.

**S3:**
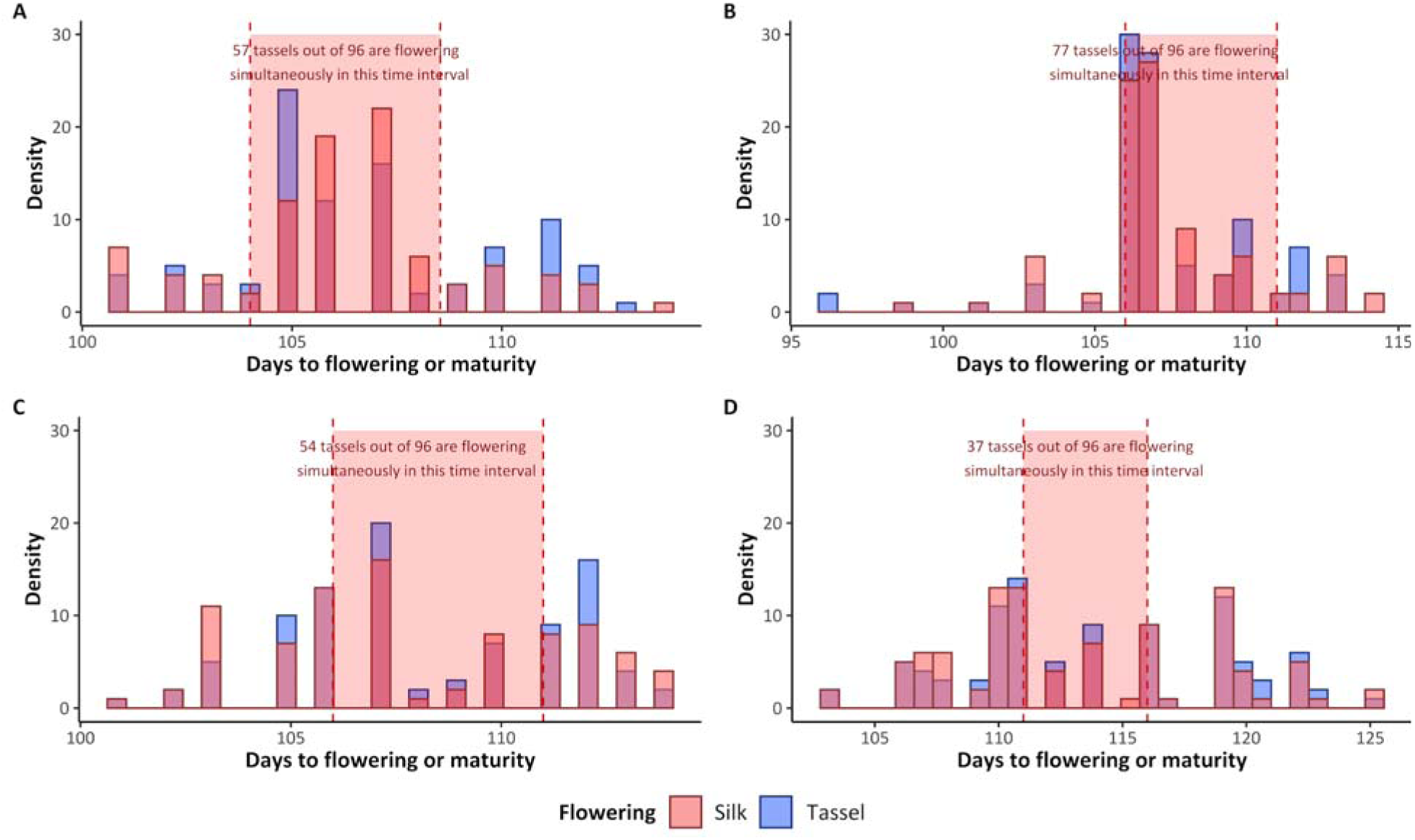
Procedure for determining the number of simultaneously flowering plants and female plants in population short 1 (A), short 2 (B) and tall 1 (C) and tall 2 (D).

**S4:**
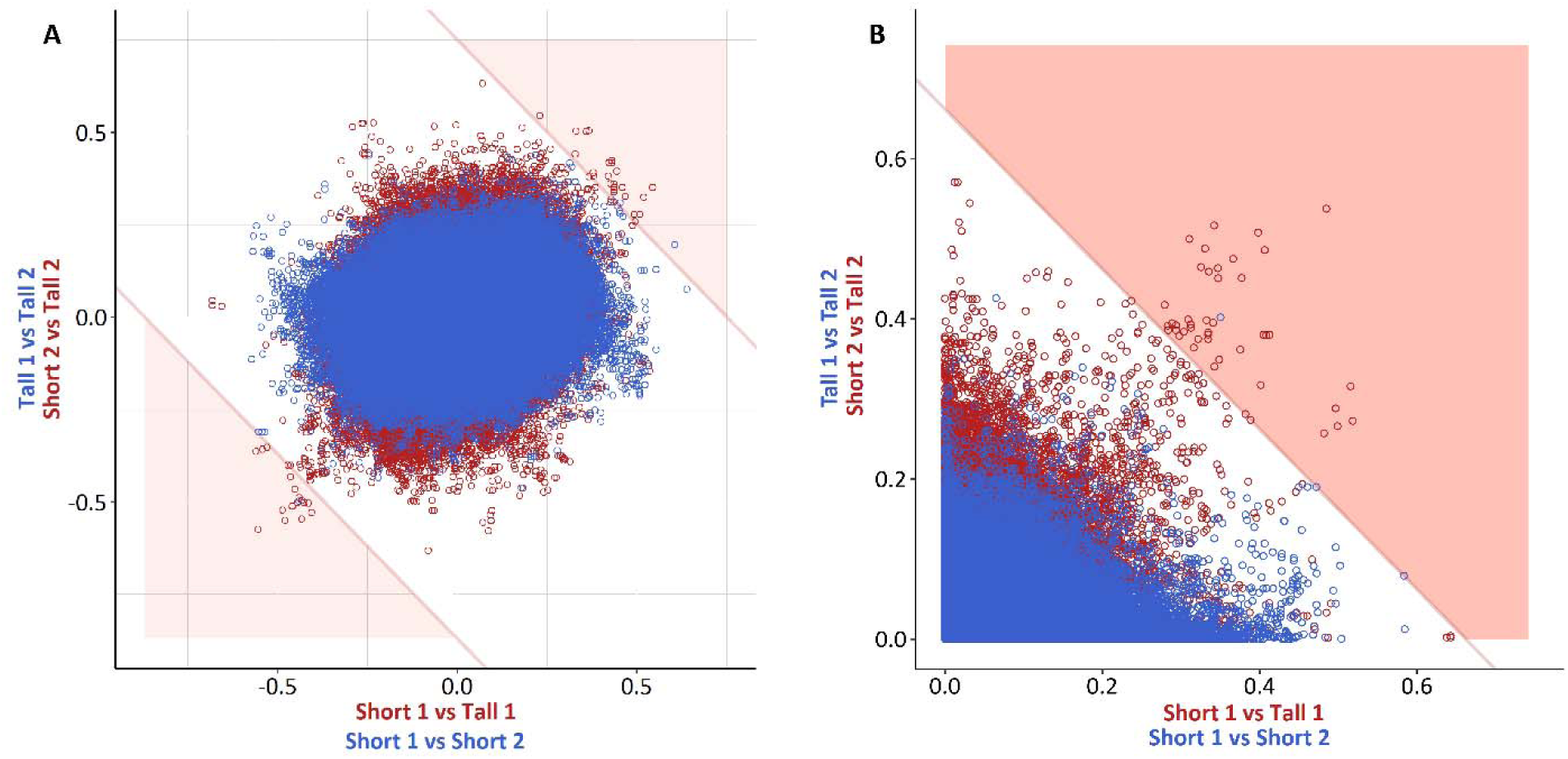
“Sauron plot” for allele frequency differences (A) and “Quarter-Sauron” plot of genetic differentiation for *F_ST_Sum* (B) observed between the subpopulations selected in the same direction (blue) and in opposite directions (red). Each dot represents one SNP. The semi-transparent red regions correspond to a false discovery rate for selection (FDRfS) <5%.

**S5:** *F_ST_Sum* values and the number of markers observed with these values computed between the subpopulations selected in the same (SD) and opposite directions (OD)

| $F_{ST}$<br>Sum | SNPs observed between<br>subpopulations selected<br>in | | FDRfS | $F_{ST}$<br>Sum | SNPs observed<br>between<br>subpopulations<br>selected in | | FDRfS | $F_{ST}$<br>Sum | SNPs observed<br>between<br>subpopulations<br>selected in | | FDRfS |
| --- | --- | --- | --- | --- | --- | --- | --- | --- | --- | --- | --- |
|  | SD | OD |  |  | SD | OD |  |  | SD | OD |  |
| 0 | 1417796 | 1421826 | 0.9972 | 0.651 | 4 | 46 | 0.0870 | 0.717 | 1 | 25 | 0.0400 |
| 0.01 | 1104313 | 1115771 | 0.9897 | 0.652 | 4 | 46 | 0.0870 | 0.718 | 1 | 24 | 0.0417 |
| 0.02 | 400204 | 415091 | 0.9641 | 0.653 | 4 | 46 | 0.0870 | 0.719 | 1 | 23 | 0.0435 |
| 0.03 | 243600 | 259984 | 0.9370 | 0.654 | 4 | 46 | 0.0870 | 0.72 | 1 | 23 | 0.0435 |
| 0.04 | 182635 | 198752 | 0.9189 | 0.655 | 4 | 45 | 0.0889 | 0.721 | 1 | 23 | 0.0435 |
| 0.05 | 142278 | 159055 | 0.8945 | 0.656 | 4 | 45 | 0.0889 | 0.722 | 1 | 23 | 0.0435 |
| 0.06 | 112868 | 128888 | 0.8757 | 0.657 | 4 | 45 | 0.0889 | 0.723 | 1 | 23 | 0.0435 |
| 0.07 | 90337 | 105548 | 0.8559 | 0.658 | 4 | 45 | 0.0889 | 0.724 | 1 | 23 | 0.0435 |
| 0.08 | 72791 | 87109 | 0.8356 | 0.659 | 4 | 45 | 0.0889 | 0.725 | 1 | 23 | 0.0435 |
| 0.09 | 58794 | 72032 | 0.8162 | 0.66 | 4 | 44 | 0.0909 | 0.726 | 1 | 23 | 0.0435 |
| 0.1 | 47919 | 60099 | 0.7973 | 0.661 | 4 | 44 | 0.0909 | 0.727 | 1 | 23 | 0.0435 |
| 0.11 | 39137 | 50155 | 0.7803 | 0.662 | 2 | 44 | 0.0455 | 0.728 | 1 | 23 | 0.0435 |
| 0.12 | 32127 | 41814 | 0.7683 | 0.663 | 1 | 42 | 0.0238 | 0.729 | 1 | 23 | 0.0435 |
| 0.13 | 26525 | 34991 | 0.7581 | 0.664 | 1 | 42 | 0.0238 | 0.73 | 1 | 23 | 0.0435 |
| 0.14 | 21905 | 29423 | 0.7445 | 0.665 | 1 | 42 | 0.0238 | 0.731 | 1 | 22 | 0.0455 |
| 0.15 | 18173 | 24779 | 0.7334 | 0.666 | 1 | 42 | 0.0238 | 0.732 | 1 | 22 | 0.0455 |
| 0.16 | 15063 | 20978 | 0.7180 | 0.667 | 1 | 42 | 0.0238 | 0.733 | 1 | 22 | 0.0455 |
| 0.17 | 12473 | 17818 | 0.7000 | 0.668 | 1 | 42 | 0.0238 | 0.734 | 1 | 22 | 0.0455 |
| 0.18 | 10368 | 15116 | 0.6859 | 0.669 | 1 | 42 | 0.0238 | 0.735 | 1 | 22 | 0.0455 |
| 0.19 | 8575 | 12813 | 0.6692 | 0.67 | 1 | 41 | 0.0244 | 0.736 | 1 | 22 | 0.0455 |
| 0.2 | 7189 | 10750 | 0.6687 | 0.671 | 1 | 41 | 0.0244 | 0.737 | 1 | 21 | 0.0476 |
| 0.21 | 6061 | 9106 | 0.6656 | 0.672 | 1 | 41 | 0.0244 | 0.738 | 1 | 20 | 0.0500 |
| 0.22 | 5126 | 7771 | 0.6596 | 0.673 | 1 | 41 | 0.0244 | 0.739 | 1 | 19 | 0.0526 |
| 0.23 | 4375 | 6602 | 0.6627 | 0.674 | 1 | 41 | 0.0244 | 0.74 | 1 | 19 | 0.0526 |
| 0.24 | 3633 | 5611 | 0.6475 | 0.675 | 1 | 40 | 0.0250 | 0.741 | 1 | 19 | 0.0526 |
| 0.25 | 3083 | 4782 | 0.6447 | 0.676 | 1 | 40 | 0.0250 | 0.742 | 1 | 19 | 0.0526 |
| 0.26 | 2579 | 4086 | 0.6312 | 0.677 | 1 | 39 | 0.0256 | 0.743 | 1 | 19 | 0.0526 |
| 0.27 | 2161 | 3477 | 0.6215 | 0.678 | 1 | 39 | 0.0256 | 0.744 | 1 | 19 | 0.0526 |
| 0.28 | 1820 | 2979 | 0.6109 | 0.679 | 1 | 39 | 0.0256 | 0.745 | 1 | 19 | 0.0526 |
| 0.29 | 1544 | 2543 | 0.6072 | 0.68 | 1 | 38 | 0.0263 | 0.746 | 1 | 19 | 0.0526 |
| 0.3 | 1304 | 2188 | 0.5960 | 0.681 | 1 | 37 | 0.0270 | 0.747 | 1 | 19 | 0.0526 |
| 0.31 | 1085 | 1865 | 0.5818 | 0.682 | 1 | 37 | 0.0270 | 0.748 | 1 | 19 | 0.0526 |
| 0.32 | 909 | 1561 | 0.5823 | 0.683 | 1 | 36 | 0.0278 | 0.749 | 1 | 19 | 0.0526 |
| 0.33 | 762 | 1336 | 0.5704 | 0.684 | 1 | 35 | 0.0286 | 0.75 | 1 | 19 | 0.0526 |
| 0.34 | 637 | 1175 | 0.5421 | 0.685 | 1 | 35 | 0.0286 | 0.751 | 1 | 19 | 0.0526 |
| 0.35 | 551 | 1034 | 0.5329 | 0.686 | 1 | 34 | 0.0294 | 0.752 | 1 | 19 | 0.0526 |
| 0.36 | 466 | 901 | 0.5172 | 0.687 | 1 | 34 | 0.0294 | 0.753 | 0 | 19 | 0.0000 |
| 0.37 | 396 | 780 | 0.5077 | 0.688 | 1 | 33 | 0.0303 | 0.754 | 0 | 19 | 0.0000 |
| 0.38 | 333 | 683 | 0.4876 | 0.689 | 1 | 33 | 0.0303 | 0.755 | 0 | 19 | 0.0000 |
| 0.39 | 283 | 605 | 0.4678 | 0.69 | 1 | 33 | 0.0303 | 0.76 | 0 | 19 | 0.0000 |
| 0.4 | 241 | 527 | 0.4573 | 0.691 | 1 | 33 | 0.0303 | 0.77 | 0 | 18 | 0.0000 |
| 0.41 | 215 | 483 | 0.4451 | 0.692 | 1 | 33 | 0.0303 | 0.78 | 0 | 18 | 0.0000 |
| 0.42 | 176 | 436 | 0.4037 | 0.693 | 1 | 33 | 0.0303 | 0.79 | 0 | 15 | 0.0000 |
| 0.43 | 153 | 391 | 0.3913 | 0.694 | 1 | 33 | 0.0303 | 0.8 | 0 | 10 | 0.0000 |
| 0.44 | 138 | 363 | 0.3802 | 0.695 | 1 | 33 | 0.0303 | 0.81 | 0 | 10 | 0.0000 |
| 0.45 | 113 | 327 | 0.3456 | 0.696 | 1 | 33 | 0.0303 | 0.82 | 0 | 7 | 0.0000 |
| 0.46 | 98 | 298 | 0.3289 | 0.697 | 1 | 31 | 0.0323 | 0.83 | 0 | 6 | 0.0000 |
| 0.47 | 89 | 267 | 0.3333 | 0.698 | 1 | 30 | 0.0333 | 0.84 | 0 | 5 | 0.0000 |
| 0.48 | 78 | 241 | 0.3237 | 0.699 | 1 | 30 | 0.0333 | 0.85 | 0 | 4 | 0.0000 |
| 0.49 | 70 | 215 | 0.3256 | 0.7 | 1 | 30 | 0.0333 | 0.86 | 0 | 3 | 0.0000 |
| 0.5 | 59 | 198 | 0.2980 | 0.701 | 1 | 30 | 0.0333 | 0.87 | 0 | 3 | 0.0000 |
| 0.51 | 51 | 188 | 0.2713 | 0.702 | 1 | 29 | 0.0345 | 0.88 | 0 | 3 | 0.0000 |
| 0.52 | 40 | 178 | 0.2247 | 0.703 | 1 | 29 | 0.0345 | 0.89 | 0 | 3 | 0.0000 |
| 0.53 | 32 | 157 | 0.2038 | 0.704 | 1 | 29 | 0.0345 | 0.9 | 0 | 2 | 0.0000 |
| 0.54 | 23 | 142 | 0.1620 | 0.705 | 1 | 29 | 0.0345 | 0.91 | 0 | 1 | 0.0000 |
| 0.55 | 18 | 134 | 0.1343 | 0.706 | 1 | 28 | 0.0357 | 0.92 | 0 | 1 | 0.0000 |
| 0.56 | 15 | 125 | 0.1200 | 0.707 | 1 | 28 | 0.0357 | 0.93 | 0 | 1 | 0.0000 |
| 0.57 | 12 | 113 | 0.1062 | 0.708 | 1 | 28 | 0.0357 | 0.94 | 0 | 1 | 0.0000 |
| 0.58 | 10 | 107 | 0.0935 | 0.709 | 1 | 27 | 0.0370 | 0.95 | 0 | 1 | 0.0000 |
| 0.59 | 9 | 99 | 0.0909 | 0.71 | 1 | 26 | 0.0385 | 0.96 | 0 | 1 | 0.0000 |
| 0.6 | 7 | 92 | 0.0761 | 0.711 | 1 | 26 | 0.0385 | 0.97 | 0 | 1 | 0.0000 |
| 0.61 | 7 | 86 | 0.0814 | 0.712 | 1 | 26 | 0.0385 | 0.98 | 0 | 1 | 0.0000 |
| 0.62 | 6 | 83 | 0.0723 | 0.713 | 1 | 26 | 0.0385 | 0.99 | 0 | 1 | 0.0000 |
| 0.63 | 6 | 72 | 0.0833 | 0.714 | 1 | 26 | 0.0385 | 1 | 0 | 1 | 0.0000 |
| 0.64 | 6 | 63 | 0.0952 | 0.715 | 1 | 26 | 0.0385 |  |  |  |  |
| 0.65 | 5 | 46 | 0.1087 | 0.716 | 1 | 25 | 0.0400 |  |  |  |  |

**S6:** Significance thresholds corresponding to other percentiles of the simulated distribution

| Percentile | Significance threshold |
| --- | --- |
| 99<sup>th</sup> percentile | 0.0952 |
| 99.9<sup>th</sup> percentile | 0.1277 |
| 99.99<sup>th</sup> percentile | 0.1555 |
| 99.999<sup>th</sup> percentile | 0.1827 |
| 99.9999<sup>th</sup> percentile | 0.2043 |

**S7:** Absolute allele frequency differences (AbsAFD) values and the number of SNPs observed with these values computed between the subpopulations selected in the same and opposite directions

| Abs AFD | SNPs observed between subpopulations selected in |  | FDRfs | Abs AFD | SNPs observed between subpopulations selected in |  | FDRfs | Abs AFD | SNPs observed between subpopulations selected in |  | FDRfs |
| --- | --- | --- | --- | --- | --- | --- | --- | --- | --- | --- | --- |
|  | the same direction (SD) | opposite directions (OD) |  |  | SD | OD |  |  | SD | OD |  |
| 0 | 4029092 | 4029092 | 1.0000 | 0.39 | 3963 | 6811 | 0.5819 | 0.78 | 6 | 57 | 0.1053 |
| 0.01 | 1349787 | 1353321 | 0.9974 | 0.4 | 3434 | 6034 | 0.5691 | 0.79 | 6 | 51 | 0.1176 |
| 0.02 | 485323 | 490866 | 0.9887 | 0.41 | 2982 | 5284 | 0.5643 | 0.80 | 6 | 46 | 0.1304 |
| 0.03 | 364245 | 371265 | 0.9811 | 0.42 | 2571 | 4667 | 0.5509 | 0.81 | 5 | 40 | 0.1250 |
| 0.04 | 312713 | 320343 | 0.9762 | 0.43 | 2187 | 4105 | 0.5328 | 0.82 | 5 | 35 | 0.1429 |
| 0.05 | 273329 | 282125 | 0.9688 | 0.44 | 1883 | 3595 | 0.5238 | 0.83 | 5 | 34 | 0.1471 |
| 0.06 | 240504 | 250277 | 0.9610 | 0.45 | 1610 | 3135 | 0.5136 | 0.84 | 5 | 28 | 0.1786 |
| 0.07 | 212146 | 223348 | 0.9498 | 0.46 | 1369 | 2759 | 0.4962 | 0.85 | 4 | 24 | 0.1667 |
| 0.08 | 187355 | 199547 | 0.9389 | 0.47 | 1179 | 2425 | 0.4862 | 0.86 | 3 | 22 | 0.1364 |
| 0.09 | 166014 | 179002 | 0.9274 | 0.48 | 976 | 2116 | 0.4612 | 0.87 | 1 | 18 | 0.0556 |
| 0.1 | 147328 | 160900 | 0.9156 | 0.49 | 820 | 1877 | 0.4369 | 0.88 | 1 | 17 | 0.0588 |
| 0.11 | 131038 | 144737 | 0.9054 | 0.5 | 679 | 1638 | 0.4145 | 0.89 | 1 | 16 | 0.0625 |
| 0.12 | 116706 | 130343 | 0.8954 | 0.51 | 551 | 1427 | 0.3861 | 0.90 | 1 | 14 | 0.0714 |
| 0.13 | 103839 | 117425 | 0.8843 | 0.52 | 445 | 1244 | 0.3577 | 0.91 | 1 | 13 | 0.0769 |
| 0.14 | 92437 | 105670 | 0.8748 | 0.53 | 364 | 1048 | 0.3473 | 0.92 | 1 | 11 | 0.0909 |
| 0.15 | 82509 | 95231 | 0.8664 | 0.54 | 310 | 893 | 0.3471 | 0.93 | 1 | 7 | 0.1429 |
| 0.16 | 73510 | 85709 | 0.8577 | 0.55 | 259 | 756 | 0.3426 | 0.94 | 0 | 6 | 0.0000 |
| 0.17 | 65650 | 77202 | 0.8504 | 0.56 | 222 | 638 | 0.3480 | 0.95 | 0 | 5 | 0.0000 |
| 0.18 | 58555 | 69483 | 0.8427 | 0.57 | 177 | 549 | 0.3224 | 0.96 | 0 | 5 | 0.0000 |
| 0.19 | 52253 | 62655 | 0.8340 | 0.58 | 144 | 471 | 0.3057 | 0.97 | 0 | 5 | 0.0000 |
| 0.2 | 46536 | 56367 | 0.8256 | 0.59 | 124 | 394 | 0.3147 | 0.98 | 0 | 3 | 0.0000 |
| 0.21 | 41382 | 50780 | 0.8149 | 0.6 | 104 | 340 | 0.3059 | 0.99 | 0 | 3 | 0.0000 |
| 0.22 | 36769 | 45743 | 0.8038 | 0.61 | 88 | 291 | 0.3024 | 1.00 | 0 | 3 | 0.0000 |
| 0.23 | 32630 | 41125 | 0.7934 | 0.62 | 74 | 258 | 0.2868 | 1.01 | 0 | 2 | 0.0000 |
| 0.24 | 29002 | 36997 | 0.7839 | 0.63 | 71 | 221 | 0.3213 | 1.02 | 0 | 2 | 0.0000 |
| 0.25 | 25762 | 33236 | 0.7751 | 0.64 | 62 | 195 | 0.3179 | 1.03 | 0 | 1 | 0.0000 |
| 0.26 | 22827 | 29793 | 0.7662 | 0.65 | 54 | 180 | 0.3000 | 1.04 | 0 | 1 | 0.0000 |
| 0.27 | 20141 | 26734 | 0.7534 | 0.66 | 46 | 169 | 0.2722 | 1.05 | 0 | 1 | 0.0000 |
| 0.28 | 17808 | 23977 | 0.7427 | 0.67 | 38 | 150 | 0.2533 | 1.06 | 0 | 1 | 0.0000 |
| 0.29 | 15679 | 21527 | 0.7283 | 0.68 | 33 | 136 | 0.2426 | 1.07 | 0 | 1 | 0.0000 |
| 0.3 | 13851 | 19268 | 0.7189 | 0.69 | 25 | 128 | 0.1953 | 1.08 | 0 | 1 | 0.0000 |
| 0.31 | 12180 | 17264 | 0.7055 | 0.7 | 21 | 112 | 0.1875 | 1.09 | 0 | 1 | 0.0000 |
| 0.32 | 10659 | 15435 | 0.6906 | 0.71 | 19 | 100 | 0.1900 | 1.10 | 0 | 1 | 0.0000 |
| 0.33 | 9312 | 13827 | 0.6735 | 0.72 | 16 | 90 | 0.1778 | 1.11 | 0 | 1 | 0.0000 |
| 0.34 | 8137 | 12323 | 0.6603 | 0.73 | 14 | 87 | 0.1609 | 1.12 | 0 | 1 | 0.0000 |
| 0.35 | 7113 | 10947 | 0.6498 | 0.74 | 10 | 78 | 0.1282 | 1.13 | 0 | 0 | 0.0000 |
| 0.36 | 6202 | 9742 | 0.6366 | 0.75 | 8 | 72 | 0.1111 | 1.14 | 0 | 0 | 0.0000 |
| 0.37 | 5338 | 8629 | 0.6186 | 0.76 | 8 | 70 | 0.1143 | 1.15 | 0 | 0 | 0.0000 |
| 0.38 | 4633 | 7675 | 0.6036 | 0.77 | 7 | 63 | 0.1111 | 1.16 | 0 | 0 | 0.0000 |

**S8:**
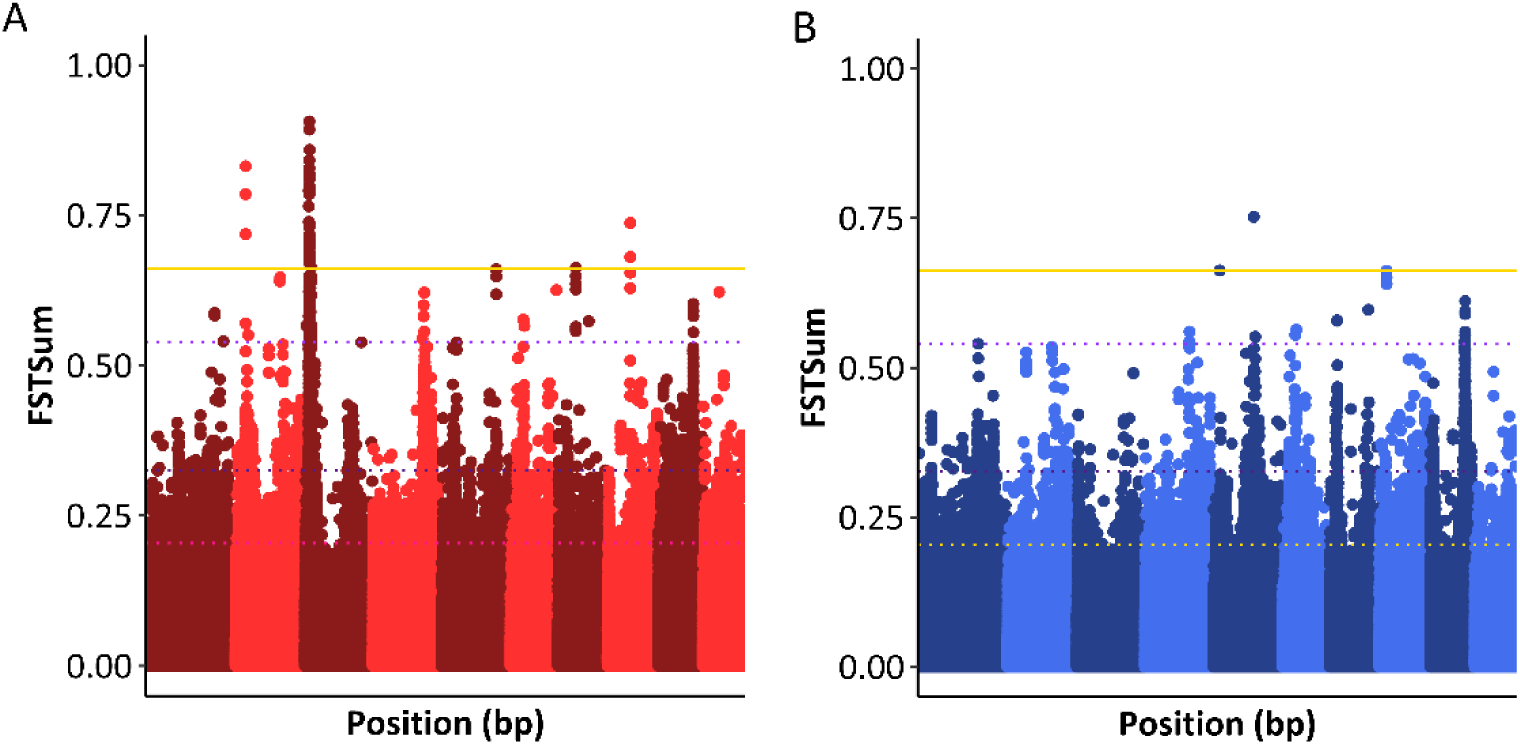
Differentiation along the entire genome expressed by *F_ST_* at each SNP marker between the subpopulations selected in opposite directions (A) and the same direction (B) with the significance thresholds based on the 99.9^th^ percentile of the empirical distribution (light purple), based on the 99.99 percentile of the empirical distribution (dark purple) and by the false discovery rate for selection (FDRfS) (yellow).

**S9:**
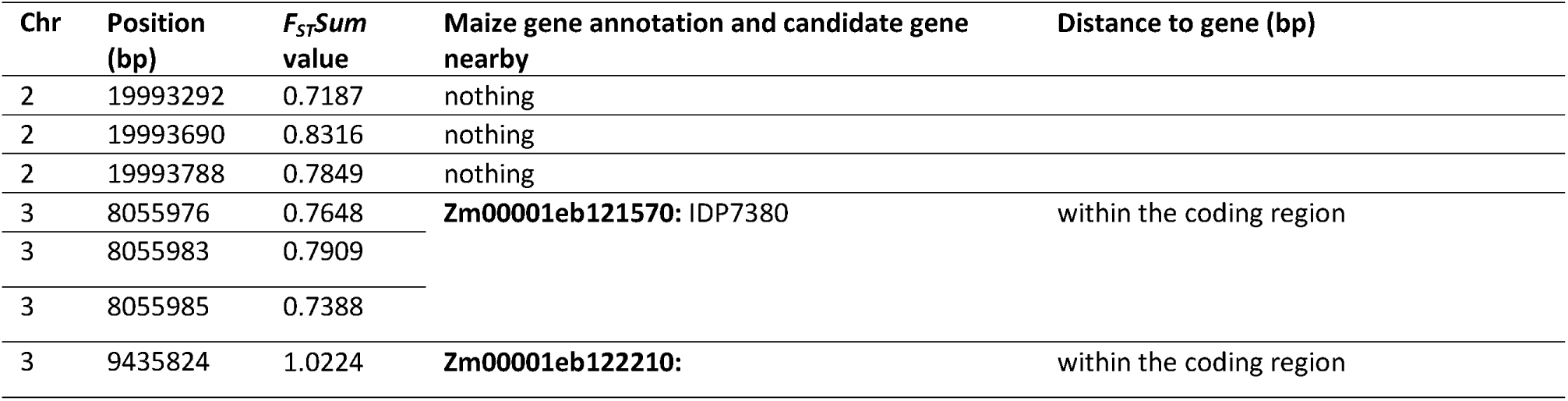
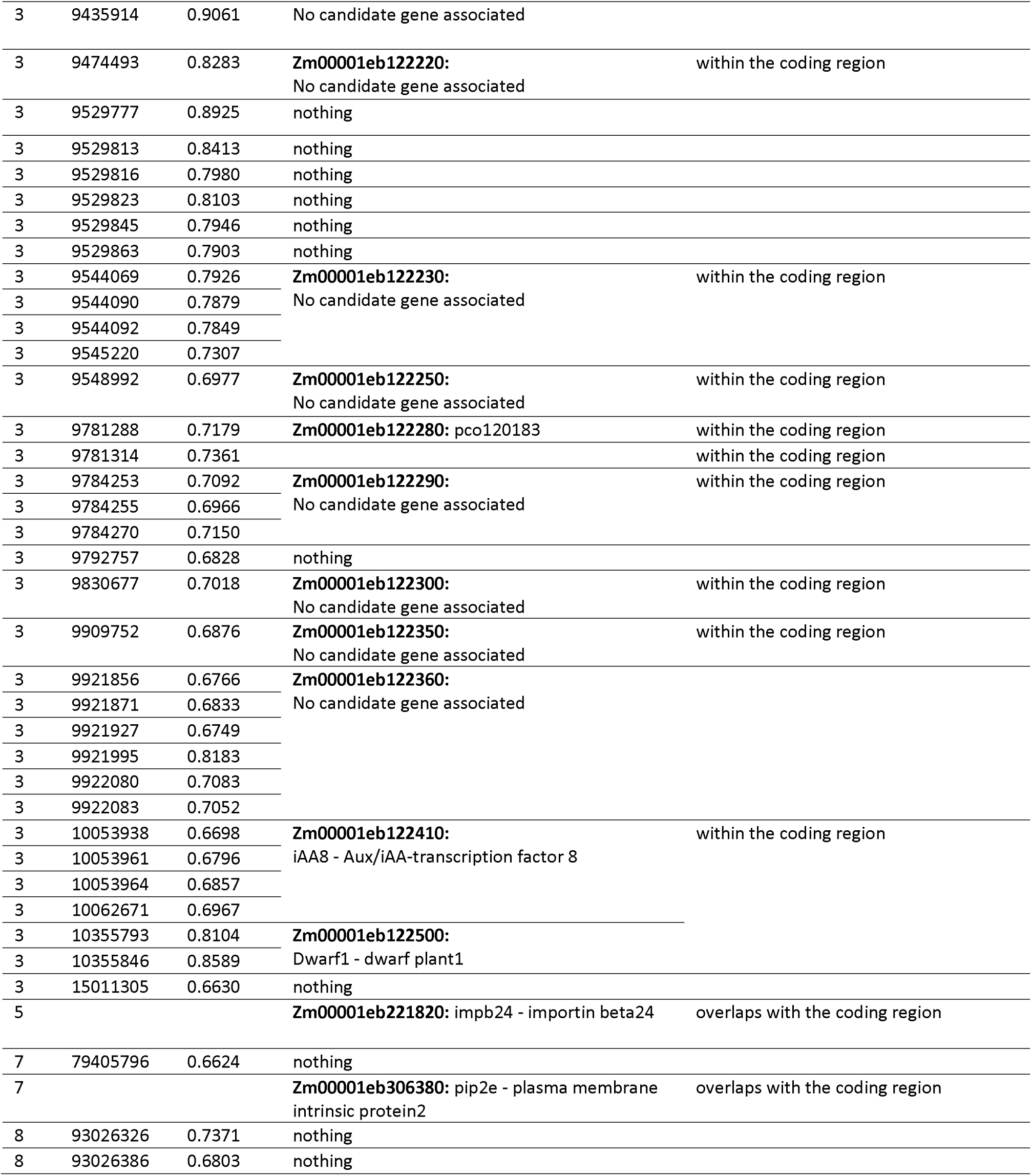
Significant SNP markers and regions with their associated gene models and candidate genes.

| Chr | Position (bp) | $F_{ST}$Sum value | Maize gene annotation and candidate gene nearby | Distance to gene (bp) |
| --- | --- | --- | --- | --- |
| 2 | 19993292 | 0.7187 | nothing |  |
| 2 | 19993690 | 0.8316 | nothing |  |
| 2 | 19993788 | 0.7849 | nothing |  |
| 3 | 8055976 | 0.7648 | **Zm00001eb121570:** IDP7380 | within the coding region |
| 3 | 8055983 | 0.7909 |  |  |
| 3 | 8055985 | 0.7388 |  |  |
| 3 | 9435824 | 1.0224 | **Zm00001eb122210:** | within the coding region |
| 3 | 9435914 | 0.9061 | No candidate gene associated |  |
| 3 | 9474493 | 0.8283 | **Zm00001eb122220:**<br>No candidate gene associated | within the coding region |
| 3 | 9529777 | 0.8925 | nothing |  |
| 3 | 9529813 | 0.8413 | nothing |  |
| 3 | 9529816 | 0.7980 | nothing |  |
| 3 | 9529823 | 0.8103 | nothing |  |
| 3 | 9529845 | 0.7946 | nothing |  |
| 3 | 9529863 | 0.7903 | nothing |  |
| 3 | 9544069 | 0.7926 | **Zm00001eb122230:**<br>No candidate gene associated | within the coding region |
| 3 | 9544090 | 0.7879 |  |  |
| 3 | 9544092 | 0.7849 |  |  |
| 3 | 9545220 | 0.7307 |  |  |
| 3 | 9548992 | 0.6977 | **Zm00001eb122250:**<br>No candidate gene associated | within the coding region |
| 3 | 9781288 | 0.7179 | **Zm00001eb122280:** pco120183 | within the coding region |
| 3 | 9781314 | 0.7361 |  | within the coding region |
| 3 | 9784253 | 0.7092 | **Zm00001eb122290:**<br>No candidate gene associated | within the coding region |
| 3 | 9784255 | 0.6966 |  |  |
| 3 | 9784270 | 0.7150 |  |  |
| 3 | 9792757 | 0.6828 | nothing |  |
| 3 | 9830677 | 0.7018 | **Zm00001eb122300:**<br>No candidate gene associated | within the coding region |
| 3 | 9909752 | 0.6876 | **Zm00001eb122350:**<br>No candidate gene associated | within the coding region |
| 3 | 9921856 | 0.6766 | **Zm00001eb122360:**<br>No candidate gene associated |  |
| 3 | 9921871 | 0.6833 |  |  |
| 3 | 9921927 | 0.6749 |  |  |
| 3 | 9921995 | 0.8183 |  |  |
| 3 | 9922080 | 0.7083 |  |  |
| 3 | 9922083 | 0.7052 |  |  |
| 3 | 10053938 | 0.6698 | **Zm00001eb122410:**<br>iAA8 - Aux/iAA-transcription factor 8 | within the coding region |
| 3 | 10053961 | 0.6796 |  |  |
| 3 | 10053964 | 0.6857 |  |  |
| 3 | 10062671 | 0.6967 |  |  |
| 3 | 10355793 | 0.8104 | **Zm00001eb122500:**<br>Dwarf1 - dwarf plant1 |  |
| 3 | 10355846 | 0.8589 |  |  |
| 3 | 15011305 | 0.6630 | nothing |  |
| 5 |  |  | **Zm00001eb221820:** impb24 - importin beta24 | overlaps with the coding region |
| 7 | 79405796 | 0.6624 | nothing |  |
| 7 |  |  | **Zm00001eb306380:** pip2e - plasma membrane intrinsic protein2 | overlaps with the coding region |
| 8 | 93026326 | 0.7371 | nothing |  |
| 8 | 93026386 | 0.6803 | nothing |  |

**S10: Detailed overview of gene models within and close to the putatively selected regions**

**S10:**
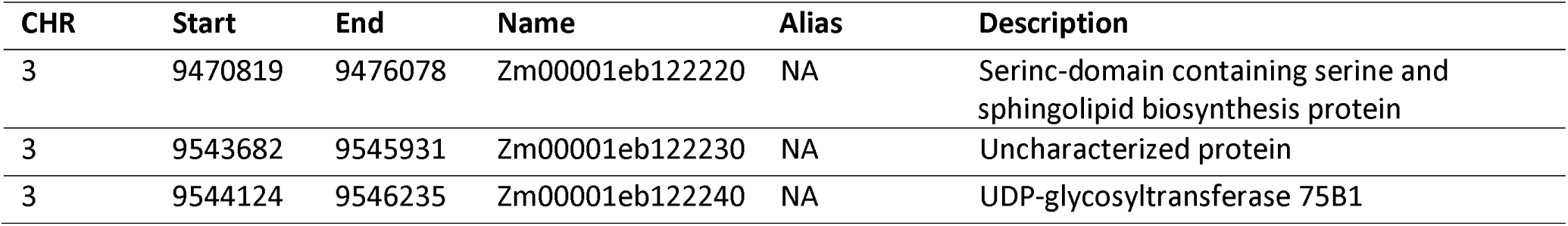
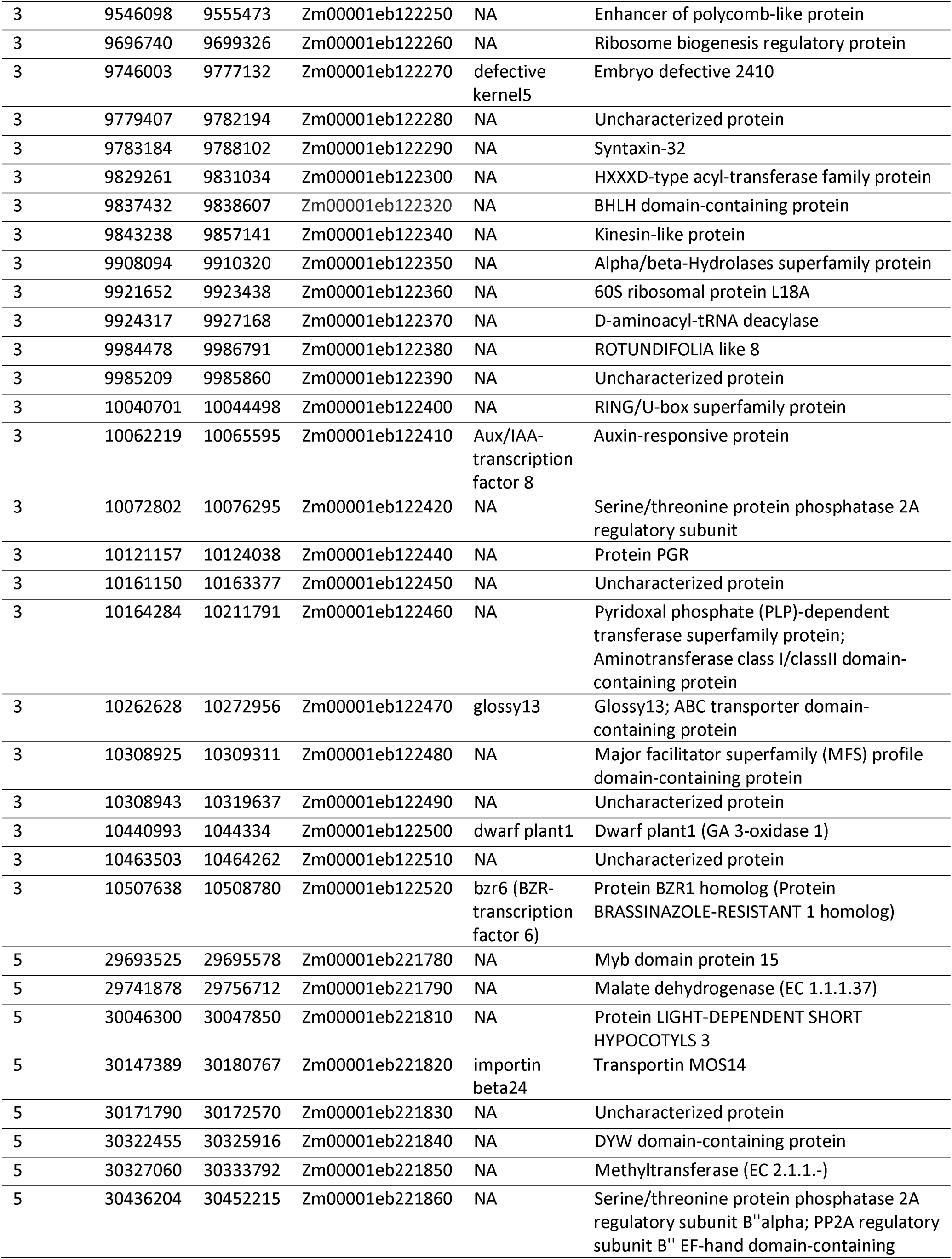
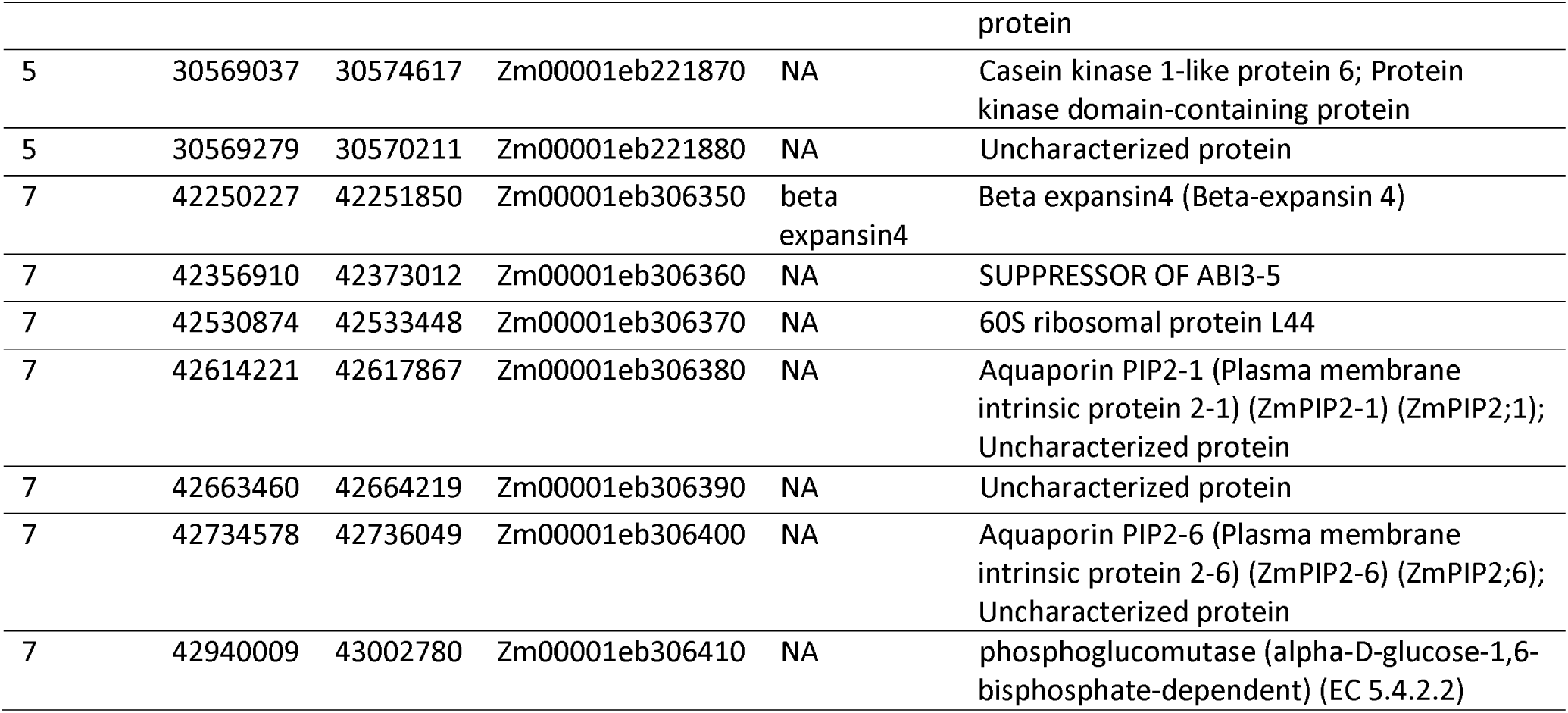
Detailed overview of gene models within and close to the putatively selected regions

**S11:**
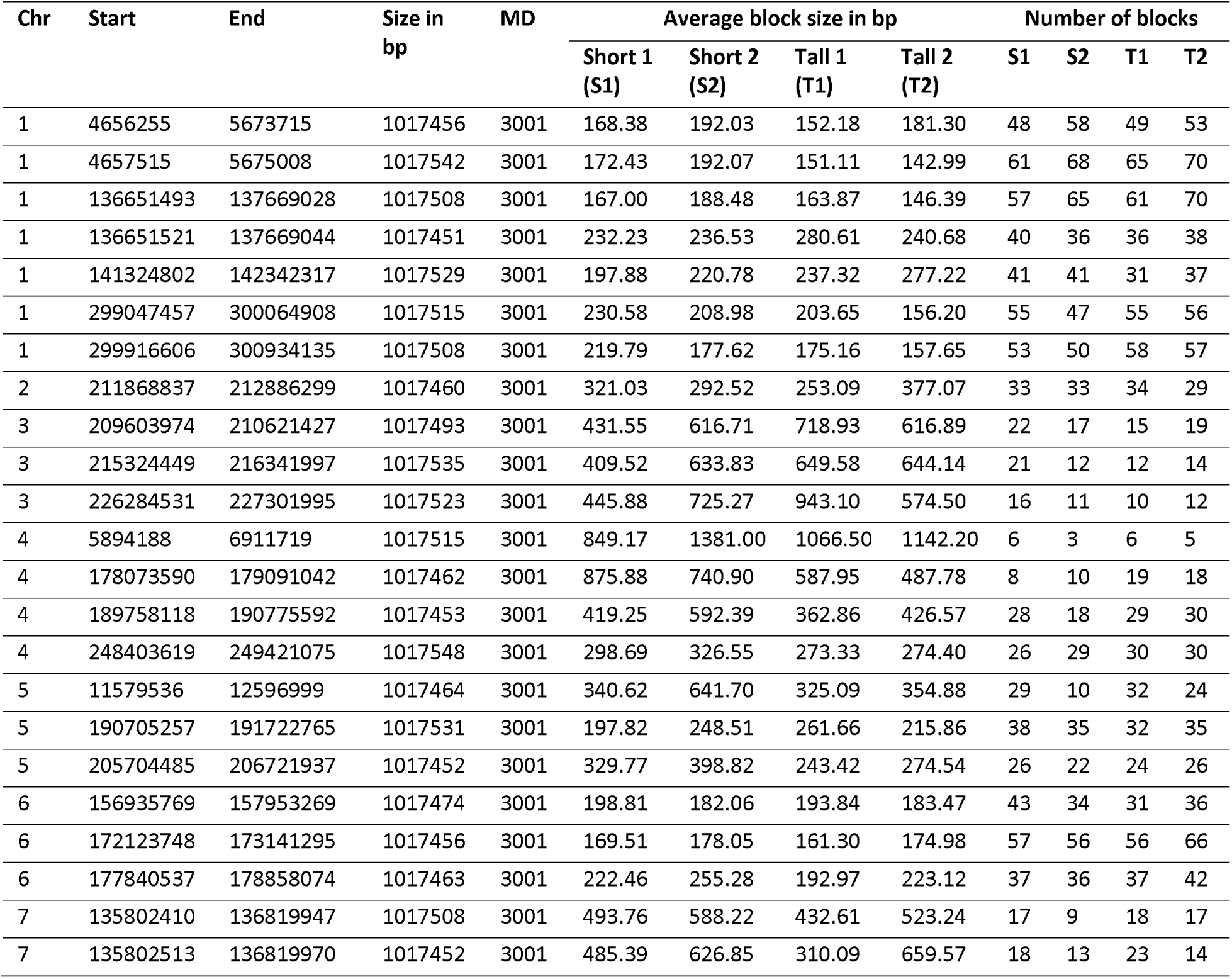
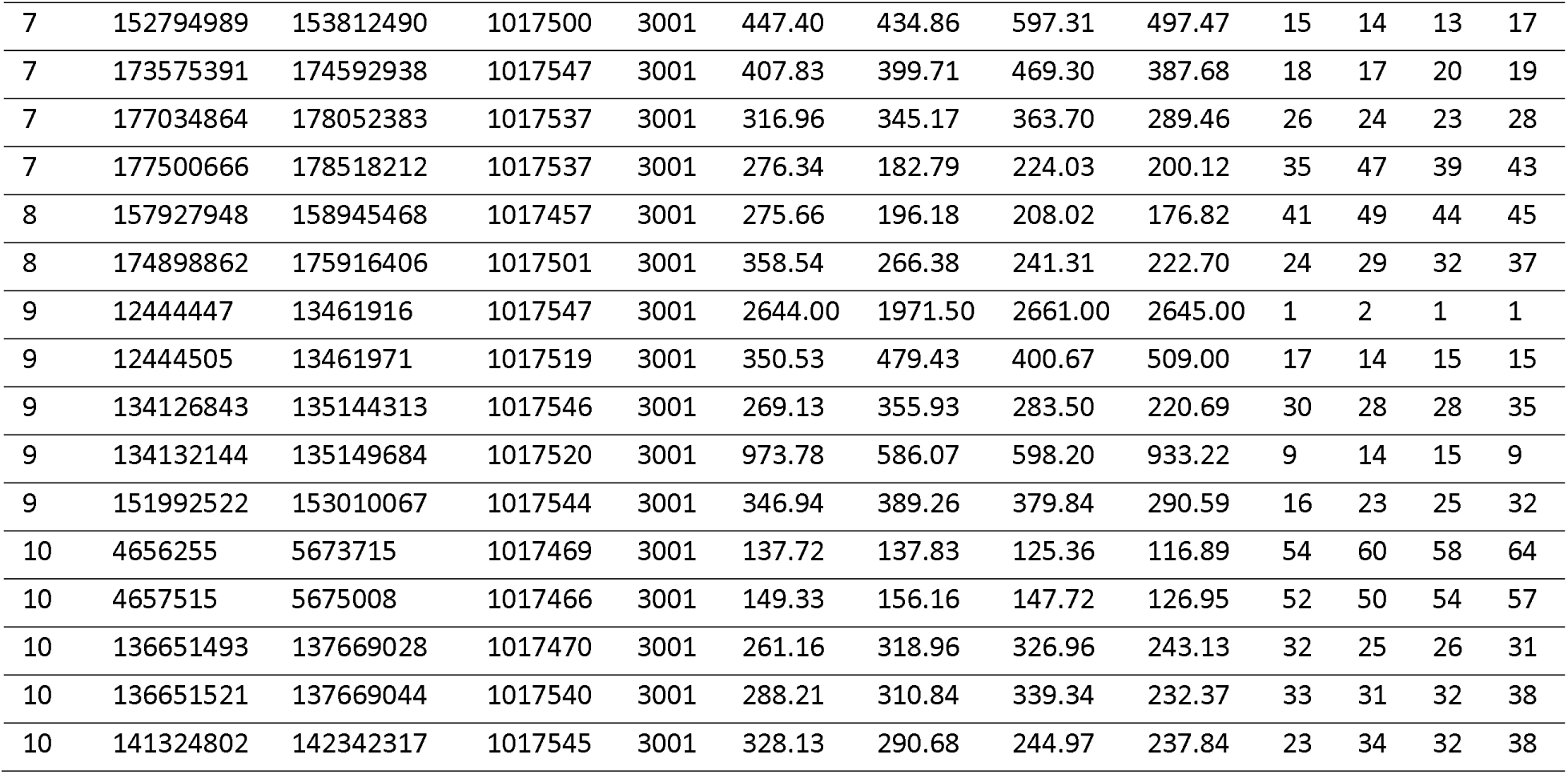
Average haplotype block size and number of blocks in the 39 randomly selected regions across the entire genome which had the same marker density (3000 markers) and the same length (1.0217 Mb) like the region on chromosome 3 from 9.411 to 10.423 Mb.

| Chr | Start | End | Size in bp | MD | Average block size in bp |  |  |  | Number of blocks |  |  |  |
| --- | --- | --- | --- | --- | --- | --- | --- | --- | --- | --- | --- | --- |
|  |  |  |  |  | Short 1 (S1) | Short 2 (S2) | Tall 1 (T1) | Tall 2 (T2) | S1 | S2 | T1 | T2 |
| 1 | 4656255 | 5673715 | 1017456 | 3001 | 168.38 | 192.03 | 152.18 | 181.30 | 48 | 58 | 49 | 53 |
| 1 | 4657515 | 5675008 | 1017542 | 3001 | 172.43 | 192.07 | 151.11 | 142.99 | 61 | 68 | 65 | 70 |
| 1 | 136651493 | 137669028 | 1017508 | 3001 | 167.00 | 188.48 | 163.87 | 146.39 | 57 | 65 | 61 | 70 |
| 1 | 136651521 | 137669044 | 1017451 | 3001 | 232.23 | 236.53 | 280.61 | 240.68 | 40 | 36 | 36 | 38 |
| 1 | 141324802 | 142342317 | 1017529 | 3001 | 197.88 | 220.78 | 237.32 | 277.22 | 41 | 41 | 31 | 37 |
| 1 | 299047457 | 300064908 | 1017515 | 3001 | 230.58 | 208.98 | 203.65 | 156.20 | 55 | 47 | 55 | 56 |
| 1 | 299916606 | 300934135 | 1017508 | 3001 | 219.79 | 177.62 | 175.16 | 157.65 | 53 | 50 | 58 | 57 |
| 2 | 211868837 | 212886299 | 1017460 | 3001 | 321.03 | 292.52 | 253.09 | 377.07 | 33 | 33 | 34 | 29 |
| 3 | 209603974 | 210621427 | 1017493 | 3001 | 431.55 | 616.71 | 718.93 | 616.89 | 22 | 17 | 15 | 19 |
| 3 | 215324449 | 216341997 | 1017535 | 3001 | 409.52 | 633.83 | 649.58 | 644.14 | 21 | 12 | 12 | 14 |
| 3 | 226284531 | 227301995 | 1017523 | 3001 | 445.88 | 725.27 | 943.10 | 574.50 | 16 | 11 | 10 | 12 |
| 4 | 5894188 | 6911719 | 1017515 | 3001 | 849.17 | 1381.00 | 1066.50 | 1142.20 | 6 | 3 | 6 | 5 |
| 4 | 178073590 | 179091042 | 1017462 | 3001 | 875.88 | 740.90 | 587.95 | 487.78 | 8 | 10 | 19 | 18 |
| 4 | 189758118 | 190775592 | 1017453 | 3001 | 419.25 | 592.39 | 362.86 | 426.57 | 28 | 18 | 29 | 30 |
| 4 | 248403619 | 249421075 | 1017548 | 3001 | 298.69 | 326.55 | 273.33 | 274.40 | 26 | 29 | 30 | 30 |
| 5 | 11579536 | 12596999 | 1017464 | 3001 | 340.62 | 641.70 | 325.09 | 354.88 | 29 | 10 | 32 | 24 |
| 5 | 190705257 | 191722765 | 1017531 | 3001 | 197.82 | 248.51 | 261.66 | 215.86 | 38 | 35 | 32 | 35 |
| 5 | 205704485 | 206721937 | 1017452 | 3001 | 329.77 | 398.82 | 243.42 | 274.54 | 26 | 22 | 24 | 26 |
| 6 | 156935769 | 157953269 | 1017474 | 3001 | 198.81 | 182.06 | 193.84 | 183.47 | 43 | 34 | 31 | 36 |
| 6 | 172123748 | 173141295 | 1017456 | 3001 | 169.51 | 178.05 | 161.30 | 174.98 | 57 | 56 | 56 | 66 |
| 6 | 177840537 | 178858074 | 1017463 | 3001 | 222.46 | 255.28 | 192.97 | 223.12 | 37 | 36 | 37 | 42 |
| 7 | 135802410 | 136819947 | 1017508 | 3001 | 493.76 | 588.22 | 432.61 | 523.24 | 17 | 9 | 18 | 17 |
| 7 | 135802513 | 136819970 | 1017452 | 3001 | 485.39 | 626.85 | 310.09 | 659.57 | 18 | 13 | 23 | 14 |
| 7 | 152794989 | 153812490 | 1017500 | 3001 | 447.40 | 434.86 | 597.31 | 497.47 | 15 | 14 | 13 | 17 |
| 7 | 173575391 | 174592938 | 1017547 | 3001 | 407.83 | 399.71 | 469.30 | 387.68 | 18 | 17 | 20 | 19 |
| 7 | 177034864 | 178052383 | 1017537 | 3001 | 316.96 | 345.17 | 363.70 | 289.46 | 26 | 24 | 23 | 28 |
| 7 | 177500666 | 178518212 | 1017537 | 3001 | 276.34 | 182.79 | 224.03 | 200.12 | 35 | 47 | 39 | 43 |
| 8 | 157927948 | 158945468 | 1017457 | 3001 | 275.66 | 196.18 | 208.02 | 176.82 | 41 | 49 | 44 | 45 |
| 8 | 174898862 | 175916406 | 1017501 | 3001 | 358.54 | 266.38 | 241.31 | 222.70 | 24 | 29 | 32 | 37 |
| 9 | 12444447 | 13461916 | 1017547 | 3001 | 2644.00 | 1971.50 | 2661.00 | 2645.00 | 1 | 2 | 1 | 1 |
| 9 | 12444505 | 13461971 | 1017519 | 3001 | 350.53 | 479.43 | 400.67 | 509.00 | 17 | 14 | 15 | 15 |
| 9 | 134126843 | 135144313 | 1017546 | 3001 | 269.13 | 355.93 | 283.50 | 220.69 | 30 | 28 | 28 | 35 |
| 9 | 134132144 | 135149684 | 1017520 | 3001 | 973.78 | 586.07 | 598.20 | 933.22 | 9 | 14 | 15 | 9 |
| 9 | 151992522 | 153010067 | 1017544 | 3001 | 346.94 | 389.26 | 379.84 | 290.59 | 16 | 23 | 25 | 32 |
| 10 | 4656255 | 5673715 | 1017469 | 3001 | 137.72 | 137.83 | 125.36 | 116.89 | 54 | 60 | 58 | 64 |
| 10 | 4657515 | 5675008 | 1017466 | 3001 | 149.33 | 156.16 | 147.72 | 126.95 | 52 | 50 | 54 | 57 |
| 10 | 136651493 | 137669028 | 1017470 | 3001 | 261.16 | 318.96 | 326.96 | 243.13 | 32 | 25 | 26 | 31 |
| 10 | 136651521 | 137669044 | 1017540 | 3001 | 288.21 | 310.84 | 339.34 | 232.37 | 33 | 31 | 32 | 38 |
| 10 | 141324802 | 142342317 | 1017545 | 3001 | 328.13 | 290.68 | 244.97 | 237.84 | 23 | 34 | 32 | 38 |

**S12:**
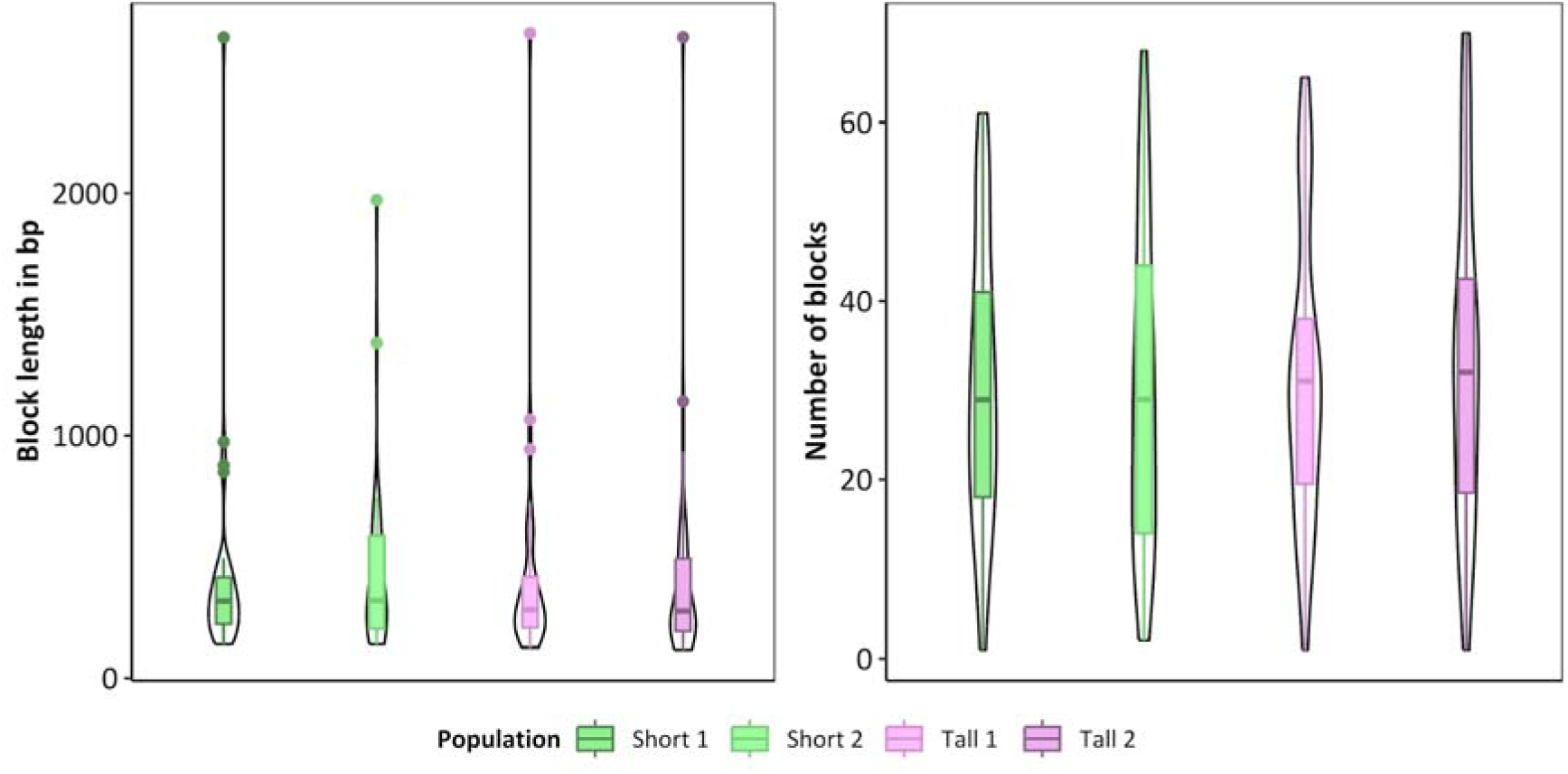
Average haplotype block size and number of blocks in the 39 randomly selected regions across the entire genome which had the same marker density (3000 markers) and the same length (1.0217 Mb) as the region on chromosome 3 from 9.411 to 10.423 Mb.

**S15:**
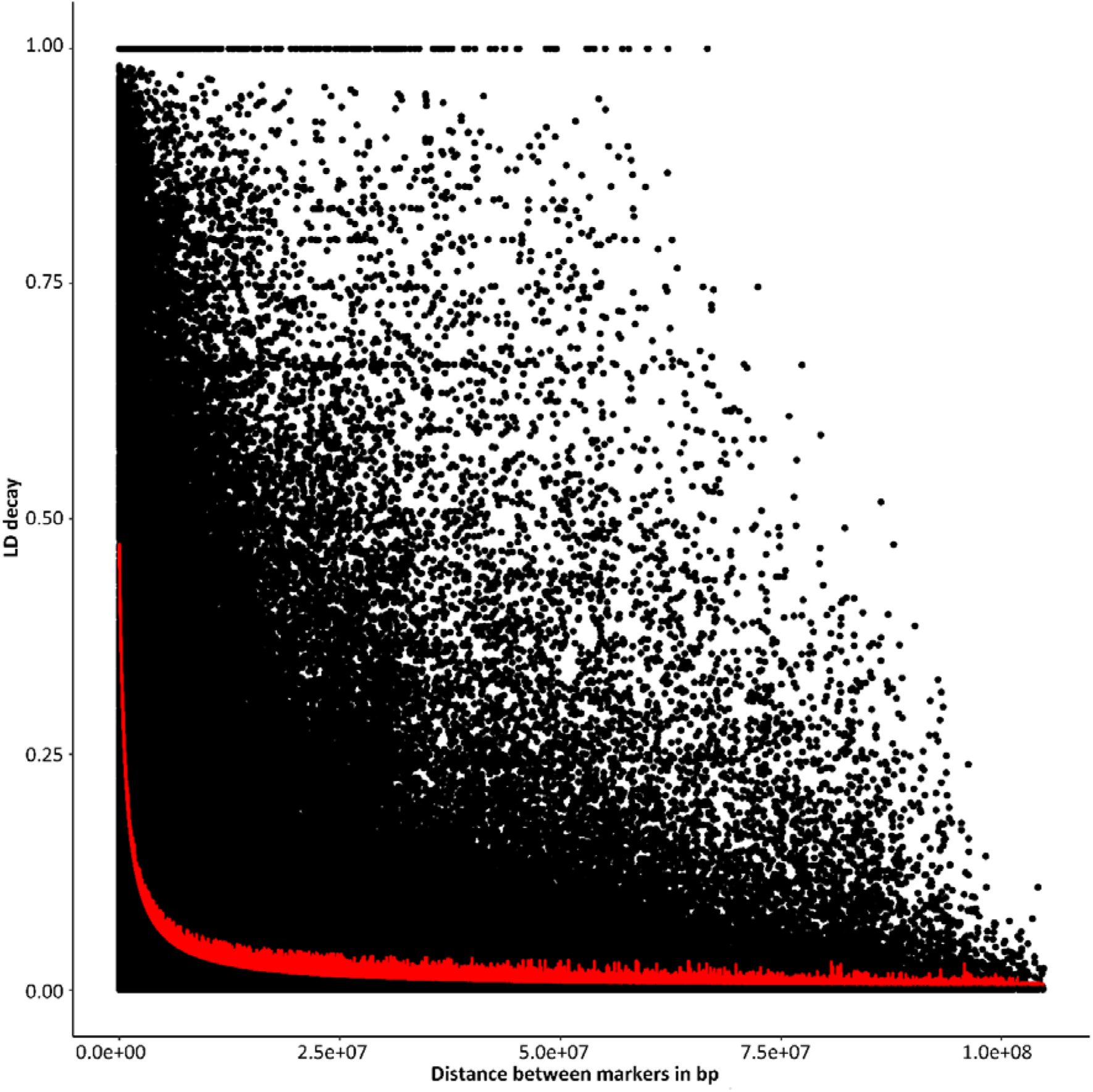
LD decay plot across the entire genome for the four subpopulations based on 10,000 randomly sampled SNPs per chromosome.

**S14:**
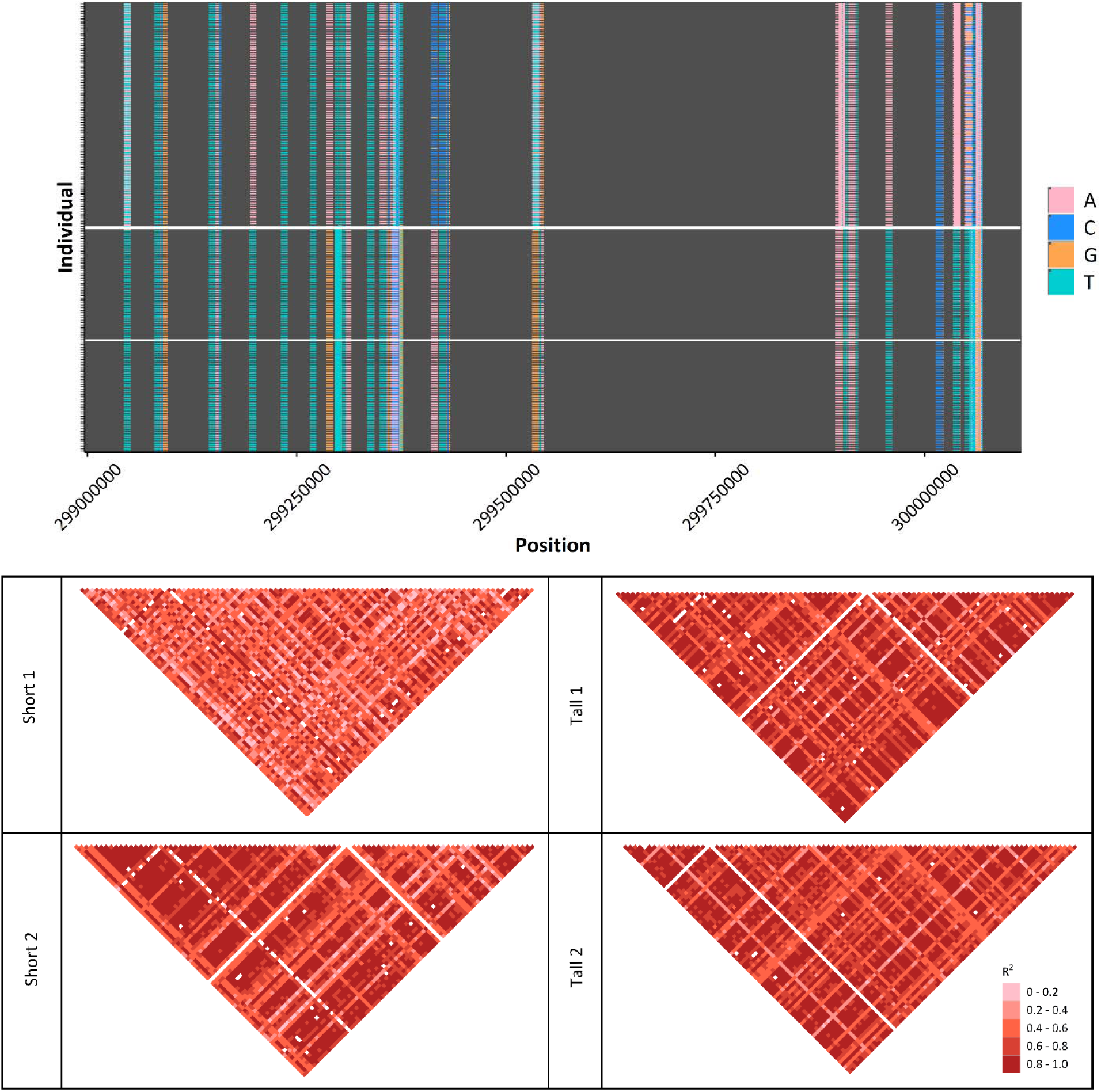
Haplotype blocks calculated in a randomly selected region on chromosome 1 from 299.05 Mb to 300.07 Mb, which had the same marker density (3000 markers) and the same length (1.0217 Mb) as the region on chromosome 3 from 9.411 to 10.423 Mb.

**S15:**
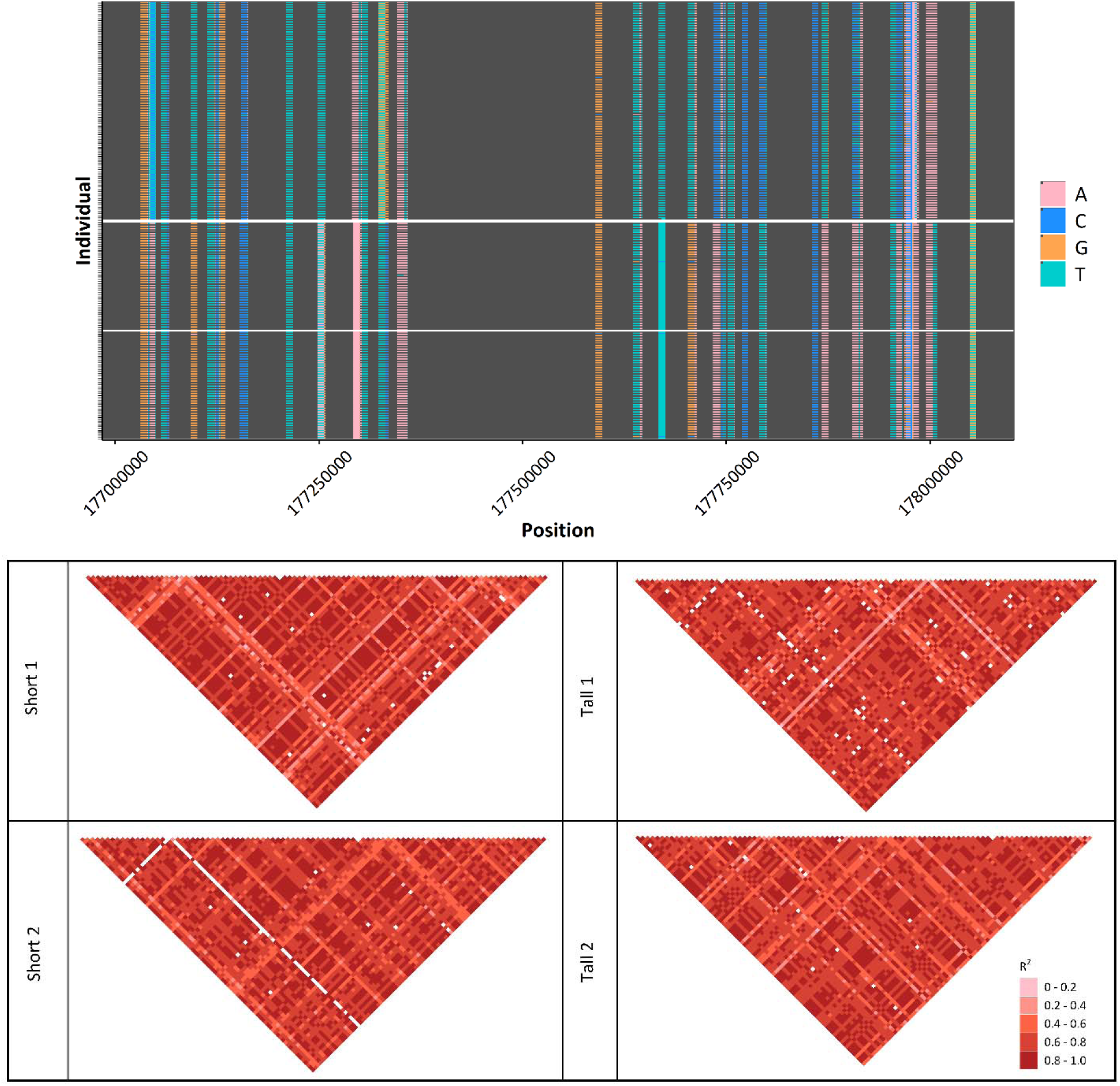
Haplotype blocks calculated in a randomly selected region on chromosome 7 from 177.04 Mb to 178.05 Mb, which had the same marker density (3000 markers) and the same length (1.0217 Mb) as the region on chromosome 3 from 9.411 to 10.423 Mb.

## References

1. Akey JM, G Zhang, K Zhang, L Jin, and MD Shriver. Interrogating a High-Density SNP Map for Signatures of Natural Selection. Genome Research. 2002;12(12): 1805–1814.

2. Akey JM. Constructing genomic maps of positive selection in humans: Where do we go from here? Genome Research. 2009;19(5): 711–722.

3. Allard RW. Principles of plant breeding. 2nd ed. New York: J. Wiley. 1999.

4. Angert AL, Bradshaw HD Jr., and Schemske DW. Using Experimental Evolution to Investigate Geographic Range Limits in Monkeyflowers. Evolution. 2008;62(10): 2660–2675.

5. Ayalew H, ME Sorrells, BF Carver, PS Baenziger, XF Ma. Selection signatures across seven decades of hard winter wheat breeding in the Great Plains of the United States. The Plant Genome. 2020;13(3).

6. Baldwin-Brown JG, A Long and KR Thornton. The power to detect quantitative trait loci using resequenced, experimentally evolved populations of diploid, sexual organisms. Molecular Biology and Evolution. 2014; 31(4): 1040–55.

7. Barghi N, R Tobler, V Nolte, C Schlötterer. Drosophila simulans: A Species with Improved Resolution in Evolve and Resequence Studies. G3 Genes|Genomes|Genetics. 2017; 7(7): 2337–2343.

8. Barghi N, R Tobler, V Nolte, AM Jakšić, F Mallard, KA Otte, M Dolezal, T Taus, R Kofler, C Schlötterer. Genetic redundancy fuels polygenic adaptation in Drosophila. PLOS Biology. 2019; 17(2): 1–31.

9. Barghi N, C Schlötterer. Shifting the paradigm in Evolve and Resequence studies: From analysis of single nucleotide polymorphisms to selected haplotype blocks. Molecular Ecology. 2019; 28: 521–524.

10. Barghi N, J Hermisson, C Schlötterer. Polygenic adaptation: a unifying framework to understand positive selection. Nature reviews. 2020; 21: 769–781.

11. Barrick JE, DS Yu, SH Yoon, H Jeong, TK Oh, D Schneider, RE Lenski, JF Kim. Genome evolution and adaptation in a long-term experiment with Escherichia coli. Nature. 2009; 461(7268): 1243–1247.

12. Beissinger TM, GJM Rosa, SM Kaeppler, D Gianola, N de Leon. Defining window-boundaries for genomic analyses using smoothing spline techniques. Genetics Selection Evolution. 2015; 47(30):1–9.

13. Beissinger TM, CN Hirsch, B Vaillancourt, S Deshpande, K Barry, CR Buell, SM Kaeppler, D Gianola, N de Leon. A Genome-Wide Scan for Evidence of Selection in a Maize Population Under Long-Term Artificial Selection for Ear Number. Genetics. 2014; 196(3): 829–840.

14. Burke MK, JP Dunham, P Shahrestani, KR Thornton, MR Rose, AD Long. Genome-wide analysis of a long-term evolution experiment with Drosophila. Nature. 2010; 467(7315): 587–590.

15. Burke MK, Liti G, Long AD. Standing Genetic Variation Drives Repeatable Experimental Evolution in Outcrossing Populations of Saccharomyces cerevisiae. Molecular Biology and Evolution. 2014;31(12): 3228–3239.

16. Bradbury PJ, ZW Zhang, DE Kroon, TM Casstevens, Y Ramdoss, ES Buckler. TASSEL: software for association mapping of complex traits in diverse samples. Bioinformatics. 2007; 23(19):2633–2635.

17. Cesconeto RJ, S Joost, CM McManus, SR Paiva, JA Cobuci, J Braccini. Landscape genomic approach to detect selection signatures in locally adapted Brazilian swine genetic groups. Ecology and Evolution. 2017; 7(22): 9544–9556.

18. Chen H, M Pelizzola, A Futschik. Haplotype based testing for a better understanding of the selective architecture. BMC Bioinformatics. 2023; 24(322):1–25.

19. Crow JF, M Kimura. An introduction to population genetics theory. 1970. Harper & Row Publishers.

20. Danecek P, JK Bonfield, J Liddle, J Marshall, V Ohan, MO Pollard, A Whitwham, T Keane, SA McCarthy, RM Davies, H Li. Twelve years of SAMtools and BCFtools. GigaScience. 2021; 10(2): giab008.

21. Doebley J, JD Wendel, JSC Smith, CW Stuber, MM Goodman. The origin of cornbelt maize: the isozyme evidence. Economic Botany. 1988; 42(1): 120 – 131.

22. Dudley JW, RJ Lambert. 100 Generations of Selection for Oil and Protein in Corn. 2003. Wiley & Sons.

23. Elshire RJ, JC Glaubitz, Q Sun, JA Poland, K Kawamoto, ES Buckler, SE Mitchell. A Robust, Simple Genotyping-by-Sequencing (GBS) Approach for High Diversity Species. PLoS ONE. 2011; 6(5), p. e19379.

24. Firman RC, F Garcia-Gonzalez, E Thyer, S Wheeler, Z Yamin, M Yuan, LW Simmons. Evolutionary change in testes tissue composition among experimental populations of house mice. Evolution. 2015; 69(3): 848–855.

25. Franssen SU, NH Barton, C Schlötterer. Reconstruction of Haplotype-Blocks Selected during Experimental Evolution. Molecular Biology and Evolution. 2016; 34(1): 174–184.

26. Galli M, Q Liua, B L. Moss, S Malcomber, W Lia, C Gainese, S Federicia, J Roshkovana, R Meeley, JL Nemhauserb, A Gallavottia. Auxin signaling modules regulate maize inflorescence architecture. Proceedings of the National Academy of Sciences. 2015; 112(43): 13372–13377.

27. Gyawali A, V Shrestha, KE Guill, S Flint-Garcia, TM Beissinger. Single-plant GWAS coupled with bulk segregant analysis allows rapid identification and corroboration of plant-height candidate SNPs. BMC Plant Biology. 2019; 19(1):412.

28. Hirsch CN, SA Flint-Garcia, TM Beissinger, SR Eichten, S Deshpande, K Barry, MD McMullen, JB Holland, ES Buckler, N Springer, CR Buell, N de Leon, SM Kaeppler. Insights into the Effects of Long-Term Artificial Selection on Seed Size in Maize. Genetics. 2014;198(1): 409–421.

29. Janick, J. Plant Breeding Reviews: Long-Term Selection: Maize, Volume 24, Part 1. 2010a. John Wiley & Sons, Inc.

30. Janick, J. (2010b) Plant Breeding Reviews, Volume 24, Part 2 Long-term Selection: Crops, Animals, and Bacteria. 2010b. John Wiley & Sons, Inc.

31. Jin, L, GB Zhang, GX Yang, JQ Dong. Identification of the Karyopherin Superfamily in Maize and Its Functional Cues in Plant Development. International Journal of Molecular Sciences. 2022; 23(14103): 1–24.

32. Kessner D, J Novembre. Power Analysis of Artificial Selection Experiments Using Efficient Whole Genome Simulation of Quantitative Traits. Genetics. Genetics, 2015; 199, 991–1005.

33. Kofler R, C Schlötterer. A Guide for the Design of Evolve and Resequencing Studies. Molecular Biology and Evolution. 2014;31(2): 474–483.

34. Krimbas CB, S Tsakas. The Genetics of Dacus oleae. V. Changes of Esterase Polymorphism in a Natural Population Following Insecticide Control-Selection or Drift? Evolution. 1971; 25(3): 454–460.

35. Kumar R, A Gyawali, GD Morrison, CA Saski, DJ Robertson, DD Cook, N Tharayil, RJ Schaefer, TM Beissinger, RS Sekhon. Genetic Architecture of Maize Rind Strength Revealed by the Analysis of Divergently Selected Populations. Plant and Cell Physiology. 2021; 62(7): 1199–1214.

36. Lai WY, V Nolte, AM Jakšić, C Schlötterer. Evolution of Phenotypic Variance Provides Insights into the Genetic Basis of Adaptation. Genome Biology and Evolution. 2024; 16(4): 1–16.

37. Li H. Aligning sequence reads, clone sequences and assembly contigs with BWA-MEM. 2013.

38. Li L, D Li, SZ Liu, XL Ma, CR Dietrich, HC Hu, GS Zhang, ZY Liu, J Zheng, GY Wang, PS Schnable. The Maize glossy13 Gene, Cloned via BSR-Seq and Seq-Walking Encodes a Putative ABC Transporter Required for the Normal Accumulation of Epicuticular Waxes. PLOS ONE 8(12): e82333.

39. Linder RA, B Zabanavar, A Majumder, H Chiao-Shyan Hoang, VG Delgado, R Tran, VT La, SW Leemans, AD Long. Adaptation in outbred sexual yeast is repeatable, polygenic, and favors rare haplotypes. Molecular Biology and Evolution, 2022; 39(12): 1–25.

40. Liu J, AR Fernie, J Yan. The Past, Present, and Future of Maize Improvement: Domestication, Genomics, and Functional Genomic Routes toward Crop Enhancement. Plant Communications. 2020;1(1),100010.

41. Long A, G Liti, A Luptak, O Tenaillon. Elucidating the molecular architecture of adaptation via evolve and resequence experiments. Nature Reviews Genetics. 2015; 16(10): 567–582.

42. Lush JL. Animal Breeding Plans. Ames, Ia., Collegiate Press. 1937.

43. Ma Y, X Ding, S Qanbari, S Weigend, Q Zhang, H Simianer. Properties of different selection signature statistics and a new strategy for combining them. Heredity. 2015; 115(5): 426–436.

44. Maize GDB. Maize genetics and genomics database. 2024. www.maizegdb.org/.

45. Mallard F, B Afonso, H Teotónio. Selection and the direction of phenotypic evolution. eLife. 2023; 12:e80993.

46. Manoli A, S Trevisan, S Quaggiotti, S Varotto. Identification and characterization of the BZR transcription factor family and its expression in response to abiotic stresses in *Zea mays* L. Plant Growth Regulation. 2018; 84: 423–436.

47. Matthes MS, NB Best, JM Robil, S Malcomber, A Gallavotti, P McSteen. Auxin EvoDevo: Conservation and Diversification of Genes Regulating Auxin Biosynthesis, Transport, and Signaling. Molecular Plant. 2019; 12(3): 298–320.

48. Mazaheri M, M Heckwolf, B Vaillancourt, JL Gage, B Burdo, S Heckwolf, K Barry, A Lipzen, C Bastos Ribeiro, TJY Kono, HF. Kaeppler, EP. Spalding, CN Hirsch, CR Buell, N de Leon, SM. Kaeppler. Genome-wide association analysis of stalk biomass and anatomical traits in maize. BMC Plant Biology. 2019; 19(45): 1–19.

49. Moose SP, JW Dudley, TR Rocheford. Maize selection passes the century mark: a unique resource for 21st century genomics. Trends in plant science. 2014; 9: 358–364.

50. Michalak P, L Kang, MF Schou, HR Garner, V Loeschcke. Genomic signatures of experimental adaptive radiation in Drosophila. Molecular Ecology. 2019; 28: 600–614.

51. Nieh SC, WS Lin, YH Hsu, GJ Shieh, BJ Kuo. The effect of flowering time and distance between pollen source and recipient on maize. GM Crops & Food. 2014; 5(4): 287–295.

52. Orozco-terwengel P, M Kapun, V Nolte, R Kofler, T Flatt, C Schlötterer. Adaptation of Drosophila to a novel laboratory environment reveals temporally heterogeneous trajectories of selected alleles. Molecular Ecology. 2012; 21: 4931–4941.

53. Otte, KA, C Schlötterer. Detecting selected haplotype blocks in evolve and resequence experiments. Molecular Ecology Resources. 2021; 21:93–109.

54. Panda C, X Li, A Wager, HY Chen, X Li. An importin-beta-like protein mediates lignin-modificationinduced dwarfism in Arabidopsis. The Plant Journal. 2020; 102: 1281–1293.

55. Parts L, FA Cubillos, J Warringer, K Jain, F Salinas, SJ Bumpstead, M Molin, A Zia, JT Simpson, MA Quail, A Moses, EJ Louis, R Durbin, G Liti. Revealing the genetic structure of a trait by sequencing a population under selection. Genome Research. 2011; 21(7): 1131–1138.

56. Payen C, AB Sunshine, GT Ong, JL Pogachar, W Zhao, MJ Dunham. High-Throughput Identification of Adaptive Mutations in Experimentally Evolved Yeast Populations. PLOS Genetics. 2016; 12(10), p. e1006339.

57. Peiffer JA, MC Romay, MA Gore, SA Flint-Garcia, Z Zhang, MJ Millard, CAC Gardner, MD McMullen, JB Holland, PJ Bradbury, ES Buckler. The genetic architecture of maize height. Genetics. 2014; 196(4): 1337–1356.

58. Palma-Vera SE, H Reyer, M Langhammer, N Reinsch, L Derezanin, J Fickel, S Qanbari, JM Weitzel, S Franzenburg, G Hemmrich-Stanisak, J Schoen. Genomic characterization of the world’s longest selection experiment in mouse reveals the complexity of polygenic traits. BMC Biology. 2022; 20(52): 1–20.

59. Phillips MA, AD Long, ZS Greenspan, LF Greer, MK Burke, B Villeponteau, KC Matsagas, CL Rizza, LD Mueller, MR Rose. Genome-wide analysis of long-term evolutionary domestication in Drosophila melanogaster. Scientific Reports. 2016; 6: 39281.

60. Phillips MA, IC Kutch, AD Long, MK Burke. Increased time sampling in an evolve-and-resequence experiment with outcrossing Saccharomyces cerevisiae reveals multiple paths of adaptive change. Molecular Ecology. 2020; 29: 4898–4912.

61. Pook, T, M Schlather, G de los Campos, M Mayer, CC Schoen, H Simianer. HaploBlocker: Creation of Subgroup-Specific Haplotype Blocks and Libraries. Genetics. 2019; 212(4): 1045–1061.

62. R Core Team. R: A Language and Environment for Statistical. 2021. https://CRAN.R-project.org/package=doParallel.

63. Ramos Báez R, RY Buckley, H Yu, Z Chen, A Gallavotti, JL Nemhauser, BL Moss. A Synthetic Approach Allows Rapid Characterization of the Maize Nuclear Auxin Response Circuit. Plant Physiology. 2020; 182(4): 1713–1722.

64. Remington DL, JM Thornsberry, Y Matsuoka, LM Wilson, SR Whitt, J Doebley, S Kresovich, MM Goodman, ES Buckler. Structure of linkage disequilibrium and phenotypic associations in the maize genome. PNAS. 2001; 98(20): 11479–11484.

65. Remolina SC, PL Chang, J Leips, SV. Nuzhdin, KA Hughes. GENOMIC BASIS OF AGING AND LIFE-HISTORY EVOLUTION IN DROSOPHILA MELANOGASTER. Evolution, 2012; 66: 3390–3403.

66. Romay MC, MJ Millard, JC Glaubitz, JA Peiffer, KL Swarts, TM Casstevens, RJ Elshire, CB Acharya, SE Mitchell, SA Flint-Garcia, MD McMullen, JB Holland, ES Buckler, CA Gardner. Comprehensive genotyping of the USA national maize inbred seed bank. Genome Biology. 2013; 14(R55).

67. Ross AJ, AR Hallauer, M Lee. Genetic analysis of traits correlated with maize ear length. Maydica. 2006; 51: 301–313.

68. Tange, O. GNU Parallel - The Command-Line Power Tool. The USENIX Magazine. 2011; 42–47.

69. Teng F, L Zhai, R Liu, W Bai, L Wang, D Huo, Y Tao, Y Zheng, Z Zhang. ZmGA3ox2, a candidate gene for a major QTL, qPH3.1, for plant height in maize. The Plant Journal. 2013; 73(3): 405–416.

70. Teotónio H, MR Rose. Variation in the reversibility of evolution. Nature. 2000; 408(6811): 463–466.

71. Tobler R, SU Franssen, R Kofler, P Orozco-terWengel, V Nolte, Joachim Hermisson, C Schlötterer. Massive Habitat-Specific Genomic Response in D. melanogaster Populations during Experimental Evolution in Hot and Cold Environments. Molecular Biology and Evolution. 2014; 31(2): 364–375.

72. Turner TL, AD Stewart, AT Fields, WR Rice, AM Tarone. Population-Based Resequencing of Experimentally Evolved Populations Reveals the Genetic Basis of Body Size Variation in Drosophila melanogaster. PLoS Genetics. 2011; 7(3): e1001336.

73. Turner TL, Miller PM. Investigating Natural Variation in Drosophila Courtship Song by the Evolve and Resequence Approach. Genetics. 2012; 191(2): 633–642.

74. Tusuubira SK, JK Kelly. Experimental evolution suggests rapid assembly of the ‘selfing syndrome’ from standing variation in *Mimulus guttatus*.Frontiers in Plant Science. 2024; 5: 1–11.

75. Scheet P, M Stephens. A fast and flexible statistical model for large-scale population genotype data: applications to inferring missing genotypes and haplotypic phase. American Journal of Human Genetics. 2006; 78(4): 629–644.

76. Schlötterer, C. How predictable is adaptation from standing genetic variation? Experimental evolution in Drosophila highlights the central role of redundancy and linkage disequilibrium. Philosphical Transactions of the Roayl Society B. 2023; 378(1877): 1–8.

77. Semagn, K, M Iqbal, N Alachiotis, A N’Diaye, C Pozniak, D Spaner. Genetic diversity and selective sweeps in historical and modern Canadian spring wheat cultivars using the 90K SNP array. Scientific Reports. 2021; 11 (23773): 1–16.

78. Silverman BW. Some Aspects of the Spline Smoothing Approach to Non-Parametric Regression Curve Fitting. Journal of the Royal Statistical Society. 1985; 47(1): 1–52

79. Shin J, S Blay, B McNeney, J Graham. “LDheatmap: An R Function for Graphical Display of Pairwise Linkage Disequilibria Between Single Nucleotide Polymorphisms.” Journal of Statistical Software. 2006; 16(3).

80. U.S. National Plant Germplasm System. Details for: PI 269743, Zea mays L. subsp. mays, Shoepeg. 2021. https://www.ars-grin.gov/.

81. Wang K, P Wu, Q Yang, D Chen, J Zhou, A Jiang, J Ma, Q Tang, W Xiao, Y Jiang, L Zhu, X Li and G Tang. Detection of Selection Signatures in Chinese Landrace and Yorkshire Pigs Based on Genotyping-by-Sequencing Data. Frontiers in Genetics. 2018; 9:119.

82. Weir BS and CC Cockerham. Estimating F-Statistics for the Analysis of Population Structure. Evolution. 1984; 38(6): 1358–1370.

83. Wickland DP, G Battu, KA Hudson, BW Diers, ME Hudson. A comparison of genotyping-by-sequencing analysis methods on low-coverage crop datasets shows advantages of a new workflow, GB-eaSy. BMC Bioinformatics. 2017; 18:586.

84. Wu YJ, ET Thorne, RE Sharp, DJ Cosgrove. Modification of Expansin Transcript Levels in the Maize Primary Root at Low Water Potentials. Plant Physiology. 2001; 126: 1471–1479.

85. Zhang, JY, S Wu, SK Boehlein, DR McCarty, GY Song, JW Walley, A Myers, AM Settles. Maize defective kernel5 is a bacterial TamB homologue required for chloroplast envelope biogenesis. Journal of Cell Biology. 2019; 218(8): 2638–2658.

